# Arabidopsis CML13 and CML14 Have Essential And Overlapping Roles In Plant Development

**DOI:** 10.1101/2023.08.30.555572

**Authors:** Kyle Symonds, Howard Teresinski, Bryan Hau, David Chiasson, Wayne A. Snedden

## Abstract

Calmodulin-like proteins (CaM-like; CML) are the largest family of calcium-binding proteins in plants, yet the functions of most CMLs are unknown. Arabidopsis CML13 and CML14 are closely related paralogs that interact with the isoleucine-glutamine (IQ) domains of myosins, IQ-domain (IQD) proteins, and CaM-activated transcription factors (CAMTAs). Here, we explored the physiological roles of CML13 and CML14 during development by using dexamethasone-inducible RNA silencing to suppress either *CML13* or *CML14* transcript levels. In the absence of inducible suppression, *CML13*- and *CML14*-RNA-interference lines were indistinguishable from WT plants throughout development. In contrast, induction of silencing treatment led to rapid increases in RNA-hairpin production that correlated with a targeted reduction in *CML13* or *CML14* transcript levels and a range of developmental and morphological effects. RNA suppression treatment did not impair the germination of *CML13- or 14*-RNA-interference lines, but these seedlings were chlorotic, displayed high mortality, and failed to achieve seedling establishment. Induced RNA suppression of mature plants led to reduced silique length, shorter roots, and rapid leaf senescence in *CML13-* and *14*-RNA-interference plants. Plants induced for RNA suppression at 2 weeks post-germination exhibited a much stronger phenotype than treatment of 3-, 4-, or 5-week-old plants. Collectively, our data indicate that both CML13 and CML14 are essential for normal development and function across a broad range of tissues and developmental stages.

**Highlight:** CML13 and CML14 are biochemically unique among the CML family and interact with proteins that possess IQ domains. Here, we show that both paralogs are essential for normal plant growth and development.

## Introduction

The importance of calcium (Ca^2+^) ions as second messengers during information processing in plants is well established (Berridge *et al*., 2000; Clapham, 2007; Luan and Wang, 2021). Complex Ca^2+^ signals are evoked by a breadth of abiotic and biotic stimuli and participate in the regulation of downstream responses corresponding to a given stimulus. Such Ca^2+^ signals are interpreted by Ca^2+^ sensor proteins, including the evolutionary-conserved and ubiquitous sensor, calmodulin (CaM) (Zhu *et al*., 2015). Ca^2+^ sensors propagate signals via direct interaction with downstream targets and are thus essential components of signaling cascades. The three main classes of Ca^2+^ sensors in plants include calmodulin (CaM) and CaM-like (CML) proteins, calcium-dependent protein kinases (CDPKs), and calcineurin B-like proteins (CBLs) that pair with CBL-interacting protein kinases (CIPKs) (Luan and Wang, 2021). These families are unique to plants and certain protists, and in Arabidopsis alone represent about a hundred putative Ca^2+^ sensors (Zhu *et al*., 2015). CDPKs and CBL-CIPK pairs serve as catalytic responders whereas CaMs and CMLs lack any catalytic activity and are thus viewed as relay sensors (Bender and Snedden, 2013; Yip Delormel and Boudsocq, 2019; Tang *et al*., 2020). Regardless, all of these Ca^2+^ sensors are thought to function as regulatory proteins, and thus their functional context is largely defined by the identity and roles of their downstream targets.

CMLs represent the largest family of Ca^2+^ sensors in plants with Arabidopsis possessing 7 CaMs and 50 CMLs (McCormack and Braam, 2003; Zhu *et al*., 2015). This impressive diversification arose early in the evolution of terrestrial plants, perhaps in response to the challenges of coping with changing environmental conditions as sessile organisms (Edel and Kudla, 2015; Edel *et al*., 2017). Indeed, the expression of many *CML*s are induced by abiotic or biotic stress (McCormack and Braam, 2003; DeFalco *et al*., 2009; Zhu *et al*., 2015). Although the majority of CMLs remain uncharacterized and their functions unknown, reports have linked several family members to important physiological processes. Some examples include roles for CML37 and CML38 in biotic and abiotic response (Vanderbeld and Snedden, 2007; Scholz *et al*., 2014; Lokdarshi *et al*., 2016), CML39 in seedling establishment and development (Bender *et al*., 2013; Midhat *et al*., 2018), CML42 in trichome development and herbivory response (Dobney *et al*., 2009; Vadassery *et al*., 2012), CML8 in pathogen response (Zhu *et al*., 2017), and CML36 in calcium homeostasis (Astegno *et al*., 2017). Although there appears to be some functional redundancy, the phylogenetic expansion of CMLs likely reflects subspecialization among family members, and the genetic studies noted above, alongside comparative biochemical studies, support this hypothesis (Dobney *et al*., 2009; Tsai *et al*., 2013; La Verde *et al*., 2018*b*).

Among the seven subfamilies of CMLs in Arabidopsis, CML13 and its close paralog CML14 (96% identity) represent the sole members of subfamily three (McCormack and Braam, 2003; Zhu *et al*., 2015). CML13 and CML14 are unique in that they do not undergo a major conformational change in response to calcium binding (Vallone *et al*., 2016; Teresinski *et al*., 2023), a hallmark structural feature of typical calcium sensors. In CaM, and other CMLs studied to date, this structural rearrangement exposes hydrophobic regions important in target recognition and binding (La Verde *et al*., 2018*a*). Moreover, public transcriptome and proteome databases indicate that both CML13 and CML14 are among the most ubiquitously expressed members of the family, suggesting they may play an important role in a variety of cell types and tissues (Teresinski *et al*., 2023).

A recent study identified CML13 and CML14 as interactors of tandem IQ (isoleucine, glutamine) domain-containing proteins including members from the IQ67-Domain cytoskeletal scaffold family (IQDs), CaM-binding transcriptional activators (CAMTAs), and myosin motor-protein families (Teresinski *et al*., 2023). In the latter case, CML13 and CML14 function as myosin light chains, and thus directly contribute to actin-myosin activity (Symonds *et al*., 2023). Collectively, the IQD, CAMTA, and myosin families cover a remarkable breadth of cell functions and are unrelated except for one structural feature, the presence of tandem IQ domains. Thus, as putative regulators of protein families that function in cytoskeletal organization, cell motility, and gene transcription, CML 13 and CML14 likely have important functions *in vivo*. However, there is a lack of genetic studies on the roles of these CMLs. Here, we explored the impact of reduced *CML13* or *CML14* transcript levels on plant morphology, growth, and development using an established, inducible RNA-suppression approach (Zhuang *et al*., 2013; Jeon *et al*., 2015; Liu and Yoder, 2016). Our data indicate that reduced transcript levels of either *CML13* or *CML14* resulted in strong, pleiotropic phenotypes including impaired seedling establishment, seedling mortality, reduced root and silique growth, and early senescence. Collectively, our findings show that both CML13 and CML14 play vital roles during various stages of plant development.

## Results

### Although 96% Identical, CML13 and CML14 Have Subtle Biochemical Differences

Differing by only eight of 148 residues (Fig 1A), CML13 and CML14 share many expression, biochemical, and functional similarities, however, there are some differences (Vallone *et al*., 2016; Teresinski *et al*., 2023). A hallmark property of typical EF-hand calcium sensors, including most CMLs, is a conformational change induced by calcium binding that exposes hydrophobic residues that are often important for interaction with downstream targets (DeFalco *et al*., 2009; La Verde *et al*., 2018*a*). Although a previous report showed that CML14 undergoes very minimal hydrophobic exposure in response to calcium, CML13 was not tested (Vallone *et al*., 2016). Thus, we used the hydrophobic probe, ANS, and compared the degree of hydrophobic exposure of recombinant CML13, CML14, and CaM7 (a conserved CaM isoform) in the presence or absence of calcium (Fig. 1B-D). As expected, a strong hydrophobic signal was seen for CaM7 in response to calcium binding (Fig. 1D). In contrast, CML14 showed almost no change in hydrophobicity (Fig. 1B), consistent with a previous report (Vallone *et al*., 2016). Our data indicate that, in general, CML13 behaves similarly to CML14 although CML14 appears to have higher constitutive hydrophobic exposure and a more discernible response to magnesium in comparison to CML13 (Fig. 1B, C). Further, CML13 and CML14 had similar melting curves (i.e. denaturation profiles) in the presence of calcium, which were 54.7 +/− 0.5 and 53.1 +/− 0.2°C, respectively (Fig. 1E, F). However, the melting curves of CML13 and CML14 in the absence of calcium (EGTA) are different by approximately 4°C, 40.7 +/− 0.7°C for CML13, and 36.5 +/− 0.3°C for CML14, suggesting that CML13 may be slightly more heat stable.

**Fig. 1.**
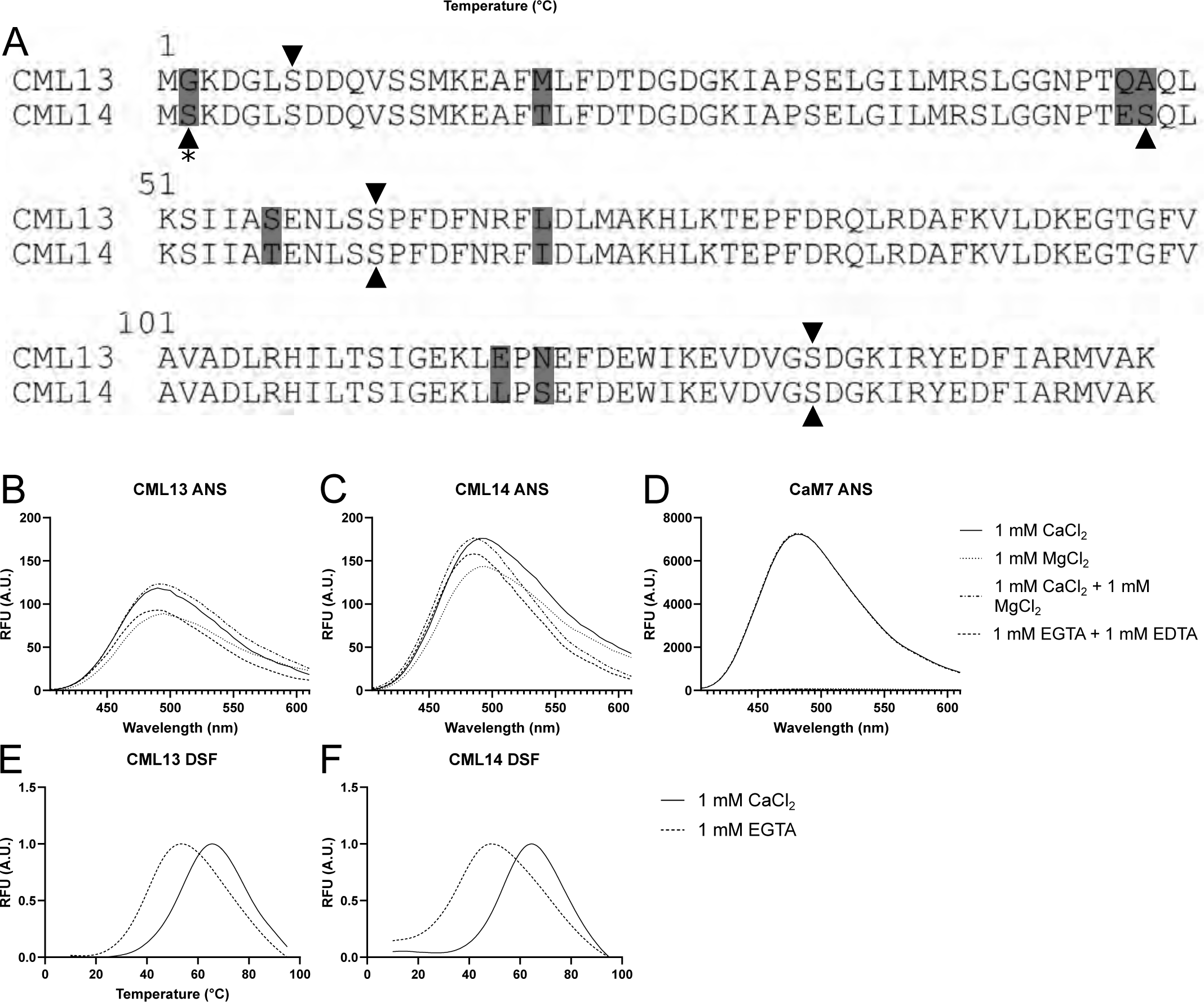
CML13 and CML14 are nearly identical proteins that differ biochemically from CaM. A) Protein sequence alignment of CML13 and CML14. Mismatched residues are highlighted in grey, phosphosites and acetylation sites, based on experimental data in public databases (PTMviewer and PhosPhAt 4.0), are represented by the arrowheads and asterisks, respectively. (B-D) Representative ANS fluorescence spectra of 20 µM CML13, CML14, and CaM7 with 1 mM CaCl_2_ (solid line), 1 mM MgCl_2_ (dotted line), 1 mM CaCl_2_ + 1 mM MgCl_2_ (dash-dot line), or 1 mM EGTA + 1 mM EDTA (dashed line). (E, F) Representative differential scanning fluorimetry (DSF) temperature spectra of (E) CML13 or (F) CML14 in the presence (solid line) or absence (dashed line) of 1 mM CaCl_2_.

### Dexamethasone-Induced RNAi-hp Reduces *CML13* and *CML14* Transcript *in vivo*

Attempts to explore the role of CML13 and CML14 by reverse genetics using stock T-DNA insertional lines from the Arabidopsis Biological Resource Center (ABRC) were hindered as databases and/or genotyping analysis indicated that the reported T-DNA insertions were unlikely to fully disrupt expression (Supplementary Fig. S1). For example, the only line (Gabi-Kat line 314F02) with a T-DNA insertion within the *CML13* ORF yielded a partial transcript that encoded the entire N-terminal lobe and most of the central linker region (Supplementary Fig. S1). CaM and some CaM-related proteins can function as single-lobe proteins (Khan and Zolman, 2010; Langelaan *et al*., 2016; Fischer *et al*., 2017). Indeed, we previously found that the isolated N-terminal lobe of CML13 can interact with the IQ region of IQD14 (Teresinski et al., 2023) and we deemed this line unsuitable for analysis in our study. Consequently, we adopted an RNA interference (RNAi-hp) strategy using the *WRKY33* intron to generate hairpins and small interfering RNAs to knock-down CML13 or CML14 (Maekawa *et al*., 2008; Zhuang *et al*., 2013; Liu and Yoder, 2016). Given that CML13 and CML14 are highly homologous at the DNA and protein levels, the hpRNAi constructs were designed to anneal to the 3’ UTR of the respective *CML13* or *CML14* mRNAs which are inherently less conserved (Fig. 2A). We opted to use the established dexamethasone (Dex)-inducible plant expression system (Zhuang *et al*., 2013; Liu and Yoder, 2016) to drive the expression of hpRNAi constructs in case the full loss-of-function of CML13 or CML14 resulted in lethality.

**Fig. 2.**
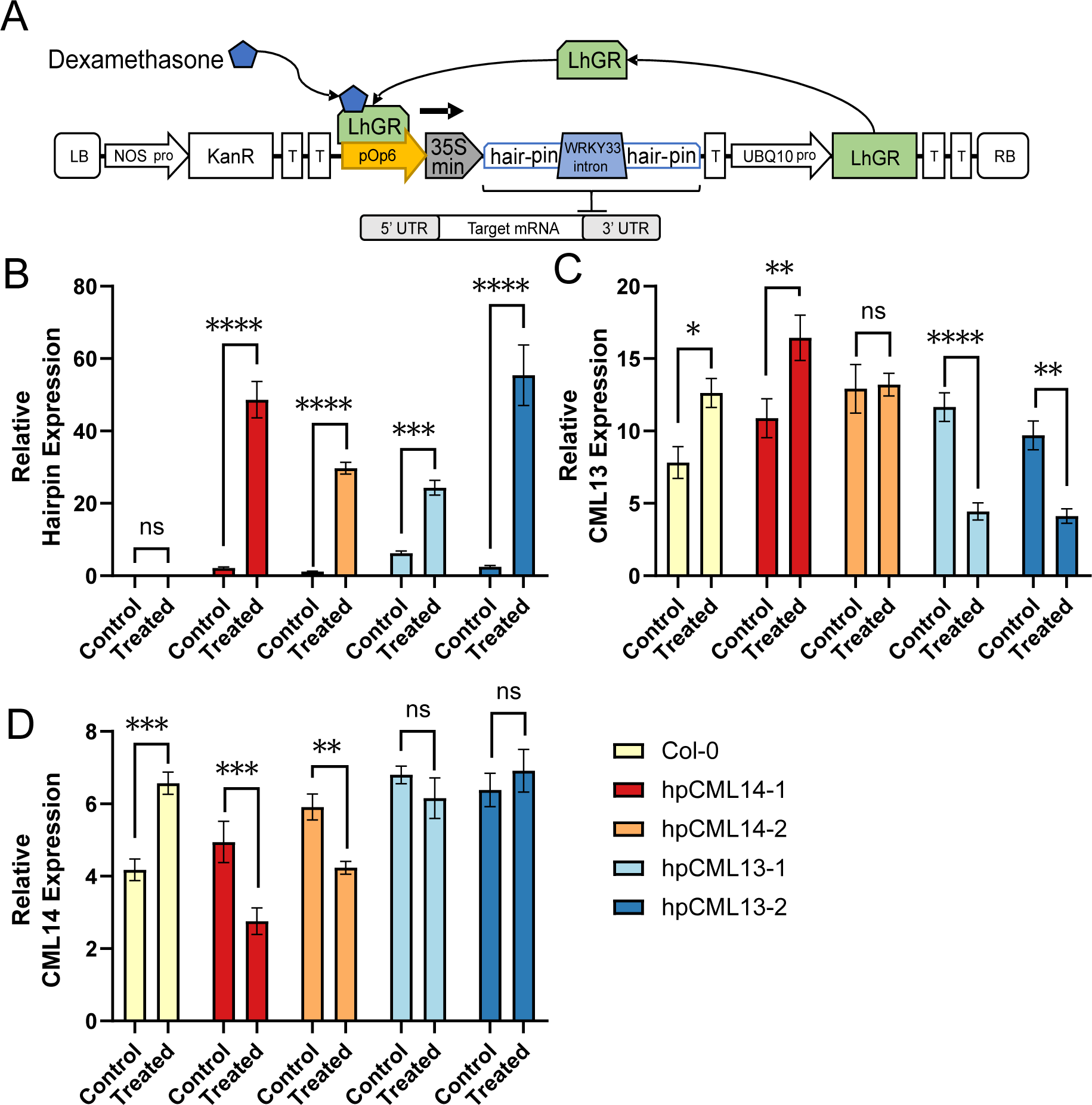
Validation of the dexamethasone (Dex)-Inducible RNA silencing of *CML13* and *CML14* transcripts. (A) Schematic diagram of the Dex-inducible RNA silencing transgene used in this study. Wild-type (Col-0) and two independent transgenic hp*CML13-* and hp*CML14* RNAi-hp lines were sown on 0.5X MS media in the presence (treated) or absence (control) of 5 µM Dex, stratified as described in the Methods, and then grown for 7 days before harvesting for mRNA extraction and analysis. (B, C, D) RT-qPCR data showing the transcript levels, normalized to *Actin2* expression, of (B) transgenic RNA hairpin transcript, (C) *CML13* transcript, and (D) *CML14* transcript in Dex-treated and control plants as indicated. All data are presented as the mean of three biological replicates with separate RNA preps of 25 - 30 plants each collected from three separate plates, and three technical reps for each RNA sample +/− SEM (2-way ANOVA with Sidak’s test for multiple comparisons, ns is not significantly different, p-value * < 0.1, ** < 0.01, *** < 0.001, **** < 0.0001).

Dex-induction of hpRNAi expression and *CML13* or *CML14* transcript reduction was monitored by RT-qPCR of 7-day-old seedlings grown on half-strength MS media with or without 5 µM Dex. In the absence of Dex, baseline expression of the RNAi-hp transcripts was extremely low, however, exposure to Dex dramatically increased RNAi-hp expression by 10-25 fold above baseline (Fig. 2B). Interestingly, in the presence of Dex, wild-type Col-0 (WT) Arabidopsis plants showed an increase in *CML13* and *CML14* transcript levels compared to untreated plants although this did not correlate with any change in discernible phenotype. This trend was also seen in the hp*CML14*-1 line which displayed elevated *CML13* transcript levels when exposed to Dex. Whether this represents a yet unknown Dex-responsive mechanism in plants is unclear. Importantly, the expression of the respective RNAi-hp caused a significant decrease of ∼30-60% of the targeted *CML13* (Fig. 2C) or *CML14* (Fig. 2D) transcripts vs control levels and we did not observe cross-suppression between these paralogs, indicating that our RNAi strategy was successful and specific.

### CML13 and CML14 are Essential for Seedling Development

To determine if CML13 and CML14 are involved in the early stages of Arabidopsis growth, WT, hp*CML13*, and hp*CML14* lines were sown onto either half-strength MS media control plates or Dex-treatment plates containing 5 µM Dex to reduce *CML13* or *CML14* transcript levels. No differential germination rate was observed for the silencing lines compared to the WT on either the control or treated medium (Fig. 3A). However, the growth and development of the hp*CML13* and hp*CML14* RNAi lines stalled at the seedling establishment stage, displaying yellow cotyledons and a reduced ability to produce true leaves (Fig. 3). Compared to WT plants or untreated RNAi-line seedlings, RNAi lines treated with Dex at 1-weeks of age showed a markedly lower proportion of green cotyledons (Fig. 3B) and reduced chlorophyll content (Fig. 3C).

**Fig. 3.**
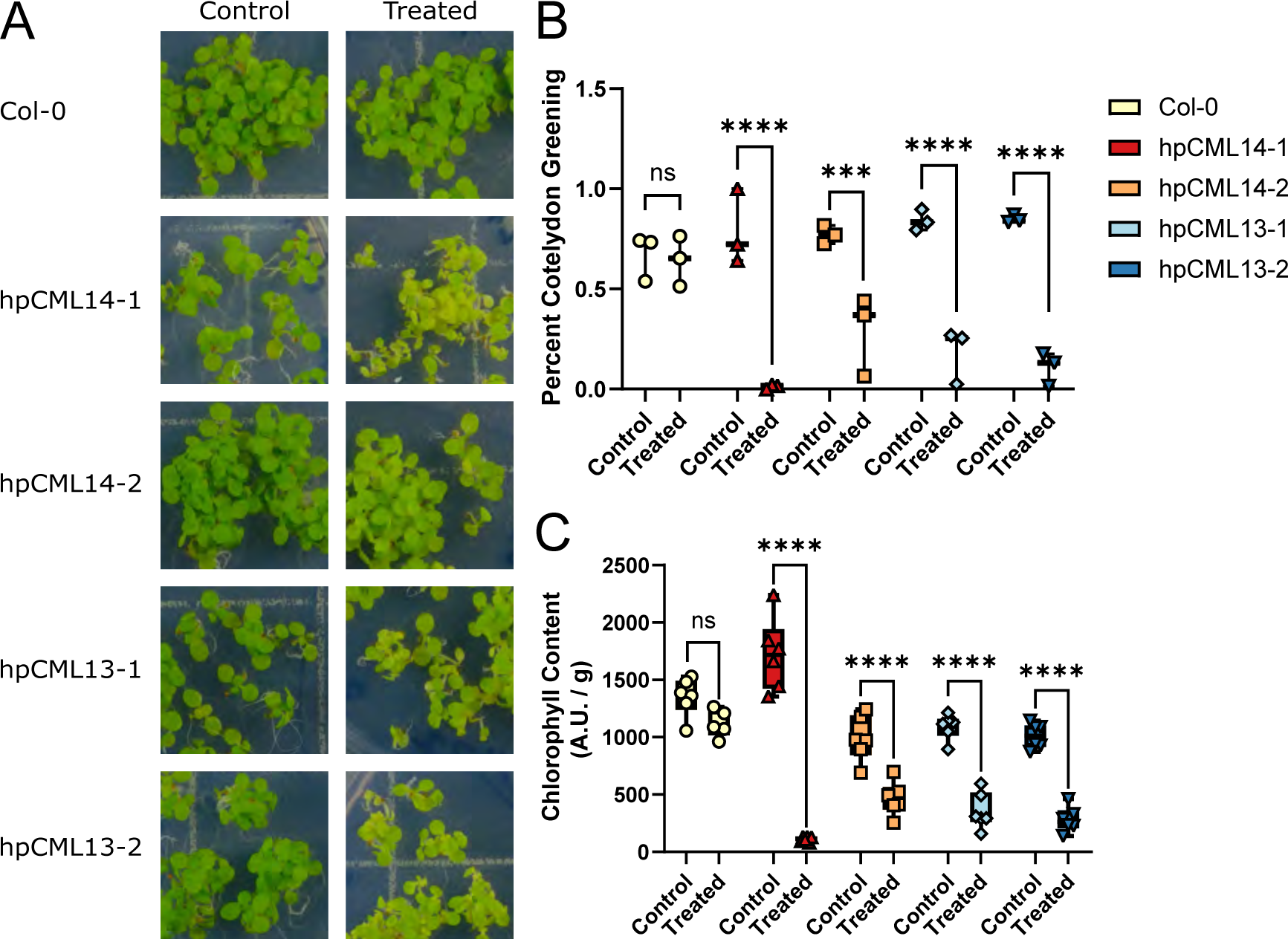
CML13 and CML14 are required for seedling establishment. (A) Representative images of wild-type (Col-0) and transgenic hp*CML* RNAi plants 10 days after stratification on control (0.5X MS) or dexamethasone (Dex)-treated (0.5X MS + 5 µM Dex) agar plates. (B) The relative number of seedlings with green cotyledons 10 days after stratification on control or Dex-treated agar plates. (C) Chlorophyll content measurements of seedlings 10 days after stratification on control or Dex-containing medium. The horizontal bar is the mean of three (B) or six (C) technical replicates each with a minimum of 50 (B) or 2 (C) seedlings (2-way ANOVA with Sidak’s test for multiple comparisons, ns is not significantly different, p-value * < .1, ** < .01, *** < .001, **** < .0001).

To explore whether CML13 or 14 are also important for development post-seedling-establishment, seedlings were grown vertically on control media for seven days before transferring to control or Dex-treated plates to measure their primary root growth rates. All plants moved to control plates grew normally (Fig. 4). Root growth of WT plants was unaffected by transfer to Dex-plates (Fig. 4A). In contrast, the hp*CML13* and hp*CML14* RNAi lines transferred onto Dex plates displayed either a reduction or cessation of primary root growth (Fig. 4B-E). Notably, the primary root growth of line hp*CML14*-1 stopped after two days on Dex-containing plates, whereas the hp*CML14*-2 line only showed statistically reduced primary root length after four days on Dex plates. This may reflect the fact that the *CML14* transcript level is not as strongly suppressed in the 14-2 line (Fig. 2D). Both of the hp*CML13* lines showed reduced primary root length after two days on Dex-containing media. Qualitatively, the Dex-treated RNAi plants appeared chlorotic, had curled leaves, and reduced rosette sizes compared to WT and untreated plants of the same lines.

**Fig. 4.**
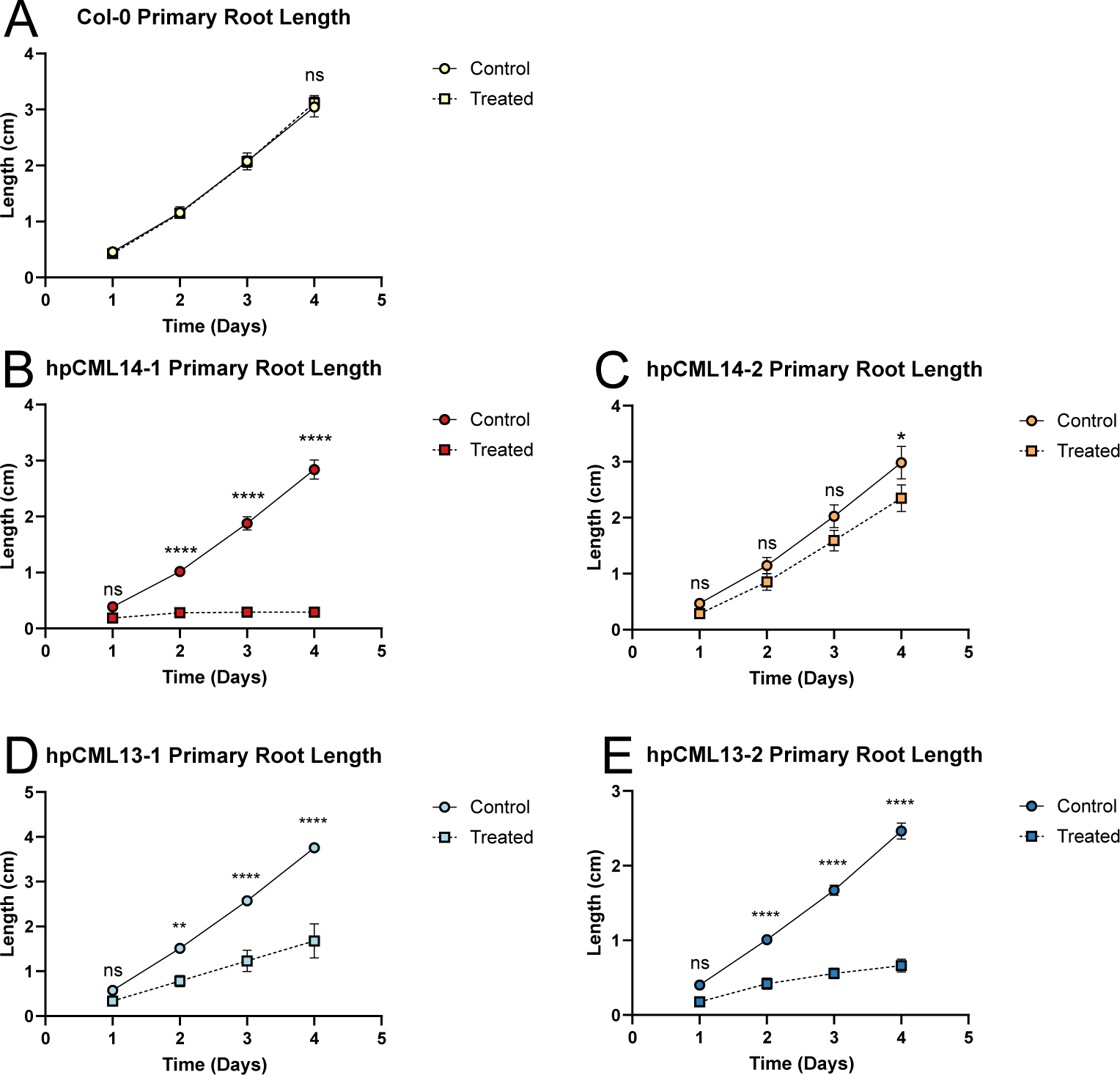
RNA-silencing of *CML13* or *CML14* inhibits primary root elongation. (A, C, E, G, and I) Representative images of 7-day-old (A) wild-type (Col-0), (C) hp*CML14*-1, (E) hp*CML14*-2, (G) hp*CML13*-1, and (I) hp*CML13*-2 seedlings transferred onto control (0.5X MS) or dexamethasone (Dex)-treated (0.5X MS + 5 µM Dex) media and grown for another 7 days after the transfer. (B, D, F, H, and J) Total primary root length of wild-type and hp*CML*RNAi transgenic lines measured per day for 4 days after transferring onto control or Dex-treated media. Points are the length of 10-16 primary roots (2-way ANOVA with Sidak’s test for multiple comparisons, ns is not significantly different, p-value * < .1, ** < .01, *** < .001, **** < .0001).

### Silencing of *CML13* or *CML14* Impairs Early Seedling Development and Induces Early Senescence

To assess the impact of silencing *CML13* or *CML14* expression during different points of vegetative and reproductive development, WT and RNAi plants were sprayed with a Dex-containing solution after one, two (Fig 5), three, four, or five weeks (Fig. 6) of growth on plates (Fig. 5A) or soil (Fig. 5C, 6). Similar to previous reports (Zhuang *et al*., 2013; Liu and Yoder, 2016), Dex application did not affect the phenotype or development of WT plants (Fig. 5, 6). One-week old hp*CML13* and hp*CML14* RNAi seedlings treated for one week with Dex on plates were chlorotic and displayed poor health (Fig. 5A). Similarly, hp*CML13* and hp*CML14* RNAi lines reared on soil for two weeks and then treated for one, two, or three weeks with Dex showed marked morphological defects, including inhibition of growth, yellowing and curling of leaves, and premature bolting and high rates of mortality (Fig. 5C). Although bolts appeared in RNAi lines, they never elongated or produced flowers and the plants continued to yellow before dying after three weeks of treatment (Fig. 5C).

**Fig. 5.**
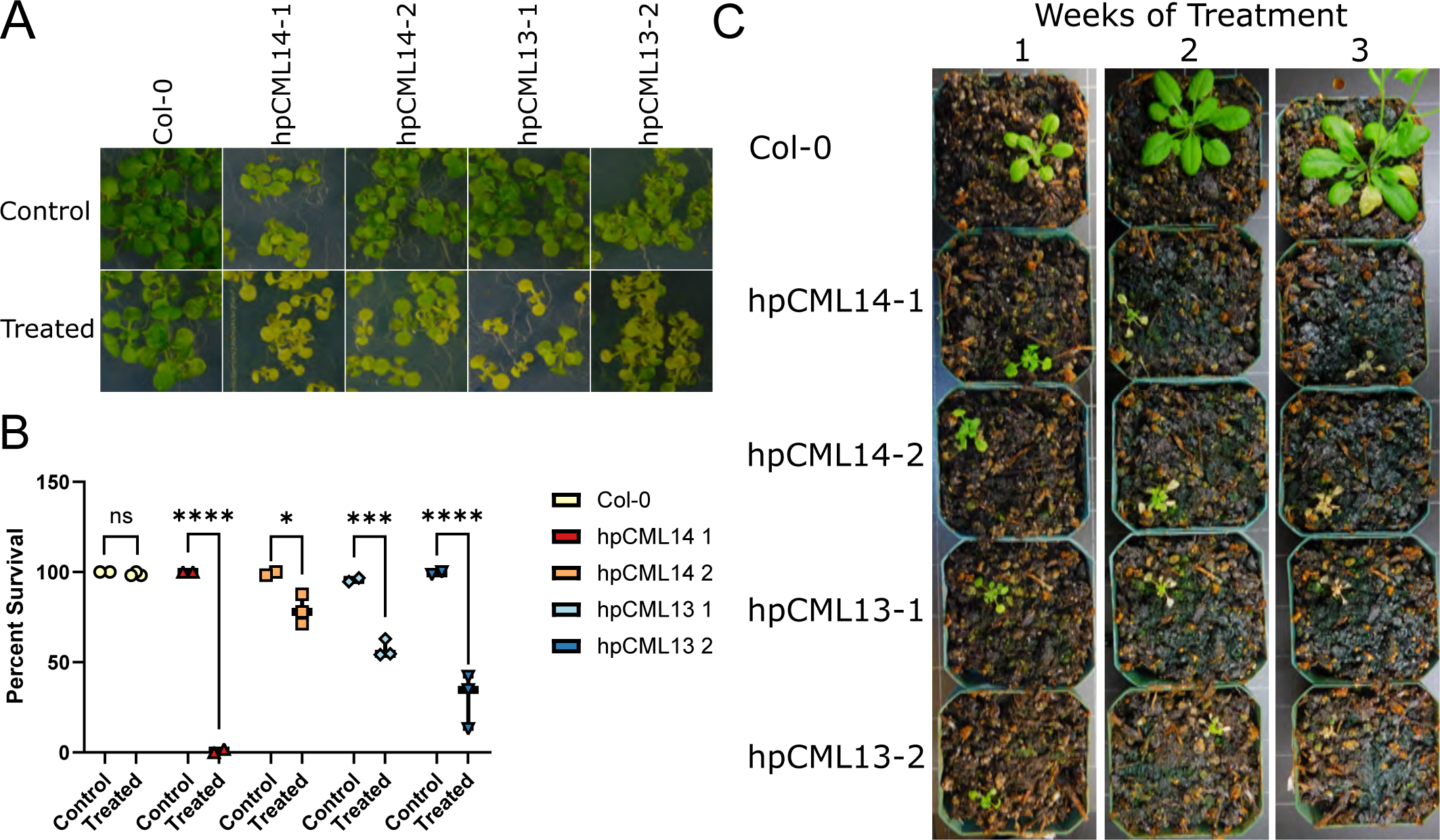
RNA-silencing of *CML13* or *CML14* early in seedling development leads to mortality. (A) Representative photos of 2-week-old Col-0 and RNAi on petri plates after one week of treatment with water or a 40 µM Dex (dexamethasone) solution. (B) Box plot showing the percent survival of either the control (water) or treatment (40 µM Dex) tabulated by the number of seedlings with yellowing/white tissue after 1 week of treatment. The horizontal bar is the mean of three technical replicates each with a minimum of 50 seedlings. (C) Representative photographs showing wild-type (Col-0), hp*CML13*-, and hp*CML14* RNAi lines that were grown for 2 weeks in soil and then sprayed with 20 µM Dex + 0.01% Silwett L-77 once every 72 hours for 1, 2, or 3 weeks, as indicated above the panel. Single plants were treated with Dex and photographs were taken over a 3-week period. Some pots have been rotated but the same plant is shown over the time course for Col-0 or each transgenic hp*CML*RNAi line. The experiment was conducted in triplicate and representative images from a single experiment are shown. (2-way ANOVA with Sidak’s test for multiple comparisons, ns is not significantly different, p-value * < .1, ** < .01, *** < .001, **** < .0001).

**Fig. 6.**
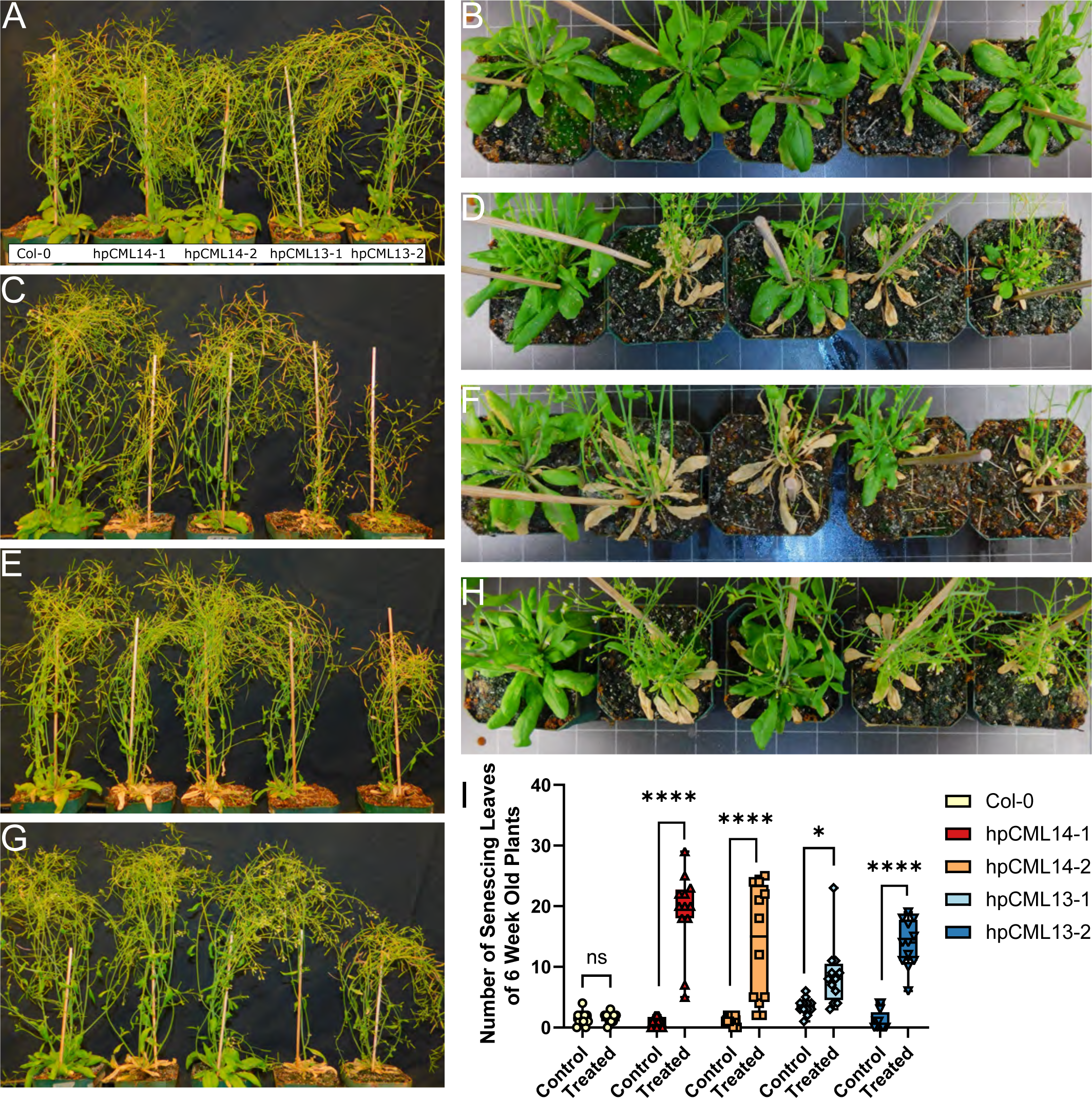
RNA-silencing *CML13* or *CML14* in mature Arabidopsis promotes early senescence. Representative side (A, C, E, G) and overhead (B, D, F, H) images of wild-type (Col-0), hp*CML13,* and hp*CML14* lines that were either (A, B) never treated with 20 µM Dex or began treatment after (C, D) 3 weeks, (E, F) 4 weeks, or (G, H) 5 weeks of age. (I) Box plot indicating the number of senescing leaves on 6-week-old plants under control conditions or treatment with 20 µM Dex + 0.01% Silwett L-77 every 72h for 4 weeks (as in panels E, F). Data points are derived from four technical replicates each with three plants from either control or Dex-treated groups each. (2-way ANOVA with Sidak’s test for multiple comparisons, ns is not significantly different, p-value * < .1, ** < .01, *** < .001, **** < .0001).

We also tested the response of mature hp*CML13* and hp*CML14* RNAi lines to Dex treatment (Fig. 6). When soil-grown three-, four-, and five-week-old WT, hp*CML13*, and hp*CML14* RNAi plants were treated with Dex up until the age of seven weeks, the hp*CML13* and hp*CML14* plants began to senesce prematurely and the rosettes were fully senesced after two weeks (Fig. 6, Supplementary Fig. S2). By seven weeks of development, the hp*CML* RNAi plants treated with Dex starting at either the three-, four-, or five-week stage, did manage to produce bolts and seeds, but displayed prematurely senesced and twisting rosette leaves (Fig. 6A-H). In general, the earlier in development that the hp*CML13* and hp*CML14* RNAi plants were treated with Dex, the more severe the observed phenotype, including reduced growth rate and shorter stature (Fig. 6C, D). More mature hp*CML* RNAi plants, sprayed with Dex from five through seven weeks of growth, showed little differences in shoot morphology compared to WT plants but had prematurely senescing rosettes similar to the RNAi plants treated earlier in development (Fig. 6G, H). We quantitatively compared the number of senescing leaves at week 6 in plants treated with Dex or control solution at week 4. Consistent with our qualitative observations (Fig. 6A-H), senescent leaves were significantly more abundant in Dex-treated *CML* hpRNAi lines compared to control plants (Fig. 6I). However, starting Dex treatment of the hpRNAi lines at these later stages of development did not have a strong effect on plant height or the number of primary and secondary bolts compared to untreated plants of the same line (Fig. S6).

### RNAi of *CML13* and *CML14* transcripts During Reproductive Phase Reduces Silique Elongation

In order to evaluate the impact of silenced *CML13* or *14* expression on silique development, WT and hp*CML* RNAi plants were treated with Dex starting at four weeks of age and continued for two weeks of treatment prior to phenotyping. Thirty mature siliques from the primary inflorescences of treated and untreated plants of each line were measured for their length. Silique length can be used as an indirect marker for reproductive success in Arabidopsis (Boyes *et al*., 2001; Bac-Molenaar *et al*., 2015). Dex-treated hp*CML13* and hp*CML14* RNAi lines, but not WT plants, showed a modest but statistically-significant reduction in silique length compared to untreated plants (Fig. 7). Interestingly, seeds from the control and treated WT or hp*CML* RNAi plants did not show any difference in germination efficiency indicating that while a reduction in *CML13* or *CML14* transcript levels impairs growth and development, seed germination efficiency may be less dependent on *CML13* or *CML14*.

**Fig. 7.**
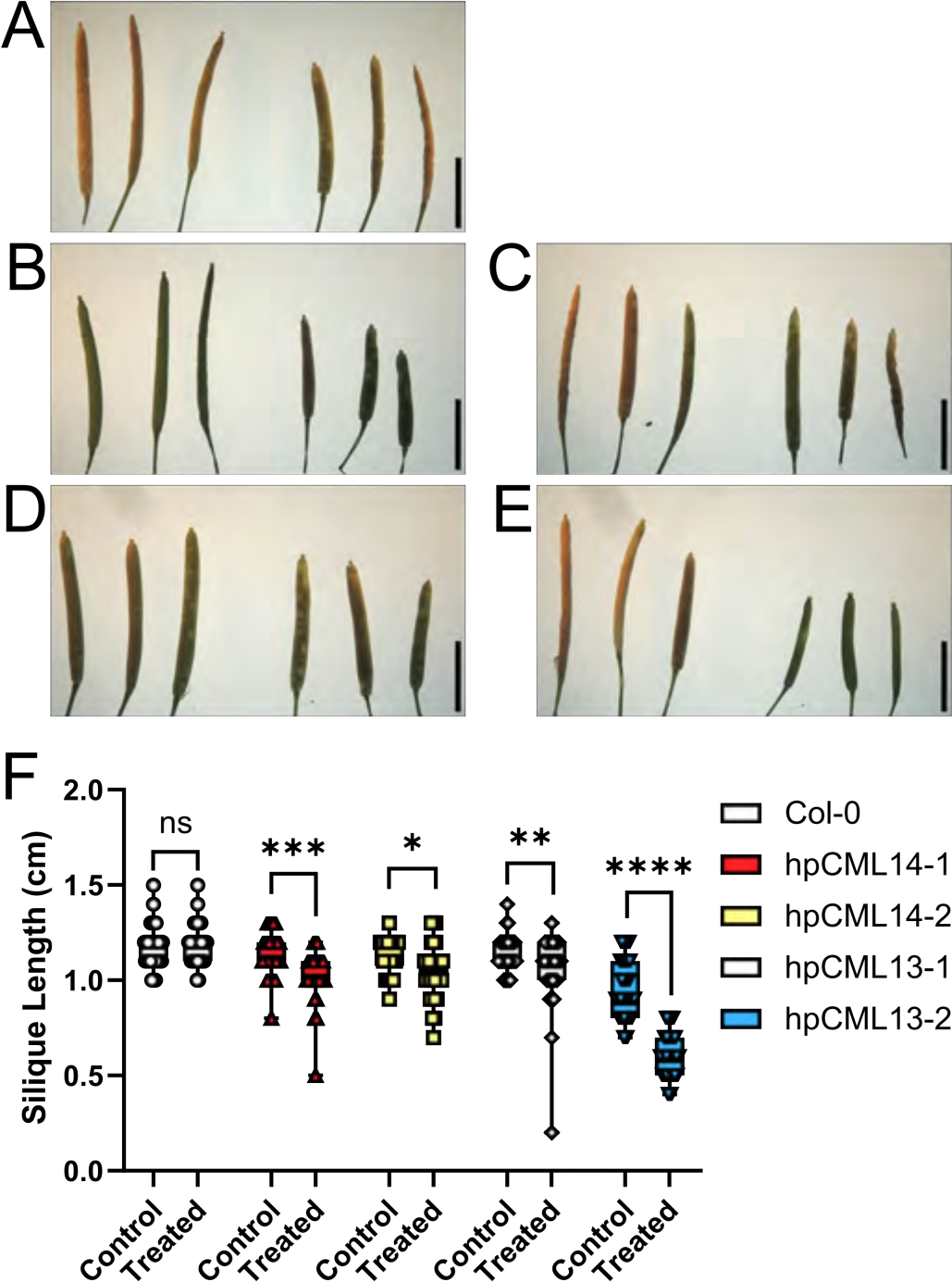
RNA-silencing of *CML13* or *CML14* reduces silique length. (A, B, C, D, and E) Representative images of mature siliques collected from (A) wild-type (Col-0), (B) hp*CML14-1*, (C) hp*CML14-2*, (D) hp*CML13-1*, and (E) hp*CML13-2* transgenic lines showing three siliques from untreated plants on the left and three siliques on the right from plants treated with 20 µM dexamethasone (Dex) every 72h starting at four weeks of age until harvesting at eight weeks of age (bar represents 5 mm). (F) Box plot of silique length for control and Dex-treated plants. Points are the lengths of 30 siliques in each control or 20 µM Dex-treated groups. (2-way ANOVA with Sidak’s test for multiple comparisons, ns is not significantly different, p-value * < .1, ** < .01, *** < .001, **** < .0001).

## Discussion

Although calcium signaling is important across a broad range of developmental processes and stress-response pathways, the roles of most plant calcium sensors remain uncharacterized (La Verde *et al*., 2018*a*). Indeed, genetic analysis has not been reported for the majority of the 50 CMLs in Arabidopsis, despite their representation as the largest family of calcium sensors. Here, using an RNA-silencing approach, we show that both CML13 and CML14 are critical for normal plant development from seedling establishment through the reproductive phase.

Among the advantages of the Dex-inducible hpRNAi system is that it facilitates the controlled spatiotemporal suppression of transcripts as transgenic lines can be progressed to homozygosity with low basal expression and good reproducibility (Zhuang *et al*., 2013; Schubert *et al*., 2022). This allows for the analysis of potentially essential genes where a complete loss-of-function would otherwise lead to lethality. Our genotyping of ABRC T-DNA lines (Supp. Fig 1) is circumstantial evidence that full loss-of-function of either *CML13* or *CML14* may indeed be lethal. The fact that very early treatment of hp*CML13* and hp*CML14* lines with Dex resulted in arrested development (Fig. 3) or lethality (Fig. 5), strongly suggests that both of these genes are essential for Arabidopsis development. The level of transcript reduction that we observed for *CML13* and *CML14* (∼30-60%, Fig. 2) is comparable to other studies on the silencing of essential genes in plants using this same technique (Zhuang *et al*., 2013; Liu and Yoder, 2016). For example, silencing the key metabolic enzyme, acetyl-CoA carboxylase, in *Medicago truncatula* roots by a Dex-induced RNA-hp resulted in a ∼60% transcript reduction and a strong root phenotype (Liu and Yoder, 2016). A similar strategy targeting an indispensable autophagy protein, SH3P2, also led to partial suppression but a strong, pleiotropic phenotype (Zhuang *et al*., 2013). However, Dex-inducible RNAi is not without limitations, including the variable time in which silencing of the target mRNA may lead to decreased protein abundance after Dex treatment. For example, monitoring YFP fluorescence intensity after differential Dex applications showed that the extent of YFP protein loss was concentration-dependent, where levels continued to decline until four days after initial Dex treatment and then remained steady and correlated with Dex concentration (Liu and Yoder, 2016). Given that RNAi inhibits translation but has no effect on extant proteins, target reduction is reliant on protein turnover. Current reports on protein turnover in Arabidopsis suggest that the median degradation rate is ∼11% of a given protein’s abundance decrease per day (Li *et al*., 2017). However, depending on the protein, tissue type, and a range of other factors, the degradation rate can vary markedly (Li *et al*., 2017). Thus, it can be difficult to predict how long after Dex treatment it will take to see changes in protein abundance and any resulting phenotype. In addition, off-target gene suppression must be considered as a possible artefact when using RNAi-based approaches. However, several lines of evidence suggest that this is not the case in our study. We targeted the unique 3’ UTRs of *CML13* or *CML14* which have very little sequence identity and we did not observe any cross-silencing between these genes (Fig. 2). Second, BLAST analysis of our RNAi constructs did not indicate any likely off-target sequences within the Arabidopsis genome. Moreover, we observed nearly identical phenotypes in Dex-treated hp*CML* plants. It is highly unlikely, given the uniqueness of the respective 3’ UTRs for *CML13* and *CML14*, that such a similar range of phenotypes would result from different off-target genes being affected. The similarity in phenotypes observed in the RNAi analysis for both *CML13* and *CML14* are consistent with their near identity at the amino-acid level, similarity in expression pattern, and overlapping profile of downstream targets (Teresinski *et al*., 2023). Thus, we are confident that the phenotypic effects we observed are directly due to the reduction of *CML13* and *CML14* transcript levels in Dex-treated plants.

While our data indicate that CML13 and CML14 are essential for seedling establishment, it is noteworthy that sowing on Dex-containing plates did not affect the germination rate of hp*CML13* or hp*CML14* lines compared to WT plants (Fig. 3). However, given the delay in suppression that ensues upon Dex treatment, noted above, it remains unclear whether CML13 or CML14 protein abundance was affected in this short period from imbibition to radical emergence, and/or whether Dex is able to penetrate the seed coat. Thus, we cannot draw conclusions as to whether CML13 and CML14 function during germination. In contrast, our data clearly indicate that CML13 or CML14 are critical during seedling establishment given that Dex treatment resulted in chlorosis, seedling arrest, cessation of primary root growth, and mortality (Fig. 3-5). Moreover, our study demonstrated that these CMLs function throughout development, as Dex application initiated at different growth stages resulted in a range of aberrant phenotypes including reduced growth, organ twisting, premature senescence, and shorter siliques (Fig. 6, 7). In general, the earlier in development that we applied Dex to hp*CML13* or hp*CML14* lines, the stronger the phenotype we observed. In contrast, the growth and development of Dex-treated WT plants were not discernibly different from untreated plants, regardless of the time of application, consistent with other studies that have used exogenous Dex application (Zhuang *et al*., 2013).

It is perhaps not surprising that suppression of *CML13* or *CML14* leads to such a strong, pleiotropic phenotype given that a recent study identified members of the CAMTA, IQD, and myosin protein families as likely CML13 and/or CML14 targets (Teresinski et al., 2023). These multigene families, comprised of 6, 33, and 17 members, respectively, function across a wide breadth of cellular processes. As key transcription factors, CAMTAs appear to be near the top of the signaling hierarchy in abiotic and biotic stress-response pathways as well as playing key roles in development (Finkler *et al*., 2007; Kim *et al*., 2009; Iqbal *et al*., 2020). Myosins are essential motor proteins that function in concert with the actin cytoskeleton to control cell and nuclear shape, cargo transport, cell division, and overall growth (Golomb *et al*., 2008; Buschmann, 2018; Haraguchi *et al*., 2019; Brueggeman *et al*., 2022). We recently provided evidence that CML13 and CML14 function as novel myosin light chains (Symonds *et al*., 2023), consistent with reports showing *in vivo* association of CML13 and CML14 with myosins (Van Leene *et al*., 2022; Teresinski *et al*., 2023). Although IQDs are not as well studied, they are thought to function as scaffolds on the cytoskeleton and have recently been implicated in cell morphology, organ and fruit shape, and cell division (Mitra *et al*., 2019; Kumari *et al*., 2021). Thus, with CAMTAs, myosins, and IQDs as putative targets, the potential number of different pathways and cellular events that may be directly or indirectly under CML13 and CML14 regulation is expansive. As such, speculation as to the mechanisms underlying the phenotype of the hp*CML* lines must be approached cautiously, as it may be due to a combination of many processes being disrupted, from changes in gene expression via CAMTAs to impacts on the cytoskeleton that alter morphology and growth rate. It is also notable that relative to other CML family members, *CML13* and *CML14* display very broad, constitutive expression and are typically among the most abundant CMLs reported in global proteomic analyses (Wang *et al*., 2015; Teresinski *et al*., 2023), suggesting they likely play housekeeping roles across tissues and throughout development.

Given the range of phenotypes observed, it is unlikely that comparative expression analysis (i.e. transcriptomics) would yield mechanistic insight into CML13/14 function, and thus moving forward, examining the effect of CML13 and CML14 on specific IQ-domain targets should be explored. Indeed, it is interesting to note that after Dex-treatment, our hpRNAi-*CML* plants appear to partially phenocopy some of the CAMTA, IQD, and myosin knockout mutants previously reported. For example, the reduced primary root length of the hp*CML13* and hp*CML14* lines is reminiscent of the reduced root length of the myosin XI-1, 2, and K triple knockout (Peremyslov *et al*., 2010). Further, the myosin XI-1, 2, K triple and XI-B, C/E double knockout mutants both display reduced silique length similar to the hp*CML13* and hp*CML14* lines after Dex treatment (Peremyslov *et al*., 2010; Madison *et al*., 2015; Tian *et al*., 2021). Moreover, both the *camta2,3* double, and *camta6* single knockouts senesce prematurely (Nie *et al*., 2012; Shkolnik *et al*., 2019), and resemble the hp*CML13 and* hp*CML14* lines (Fig. 6). In addition, when the hp*CML13 and* hp*CML14* lines were treated with Dex, they often exhibited twisted or curled rosettes (Fig. 6, Supplementary Figs S2-S5), similar to those reported for IQD5 knockout mutants (Mitra *et al*., 2019). While such correlations are noteworthy given the context of CAMTAs, IQDs, and myosins as putative CML13 and CML14 targets, further studies are needed before any mechanistic relationship can be confidently considered.

Previous studies explored the biochemical properties of CML14 (Vallone *et al*., 2016) or CML13 and CML14 (Teresinski *et al*., 2023) and concluded that these paralogs possess similar characteristics but with subtle distinctions. Our DSF and ANS fluorescence spectra of CML13 and CML14 add to that body of data and indicate that the surface-exposed hydrophobicity of these CMLs differs drastically from CaM, yet only slightly from one another (Fig. 1B-F). Changes in hydrophobicity in response to calcium binding are essential for CaM-target interaction, but whether this is also the case for CMLs is not yet known (La Verde *et al*., 2018*a*). However, we note that none of the primary sequence differences between CML13 and CML14 are in hydrophobic residues (Fig. 1A). Interestingly, phospho-proteomic databases indicate that, among those eight residues that distinguish CML13 and CML14, at least two of these positions (Ser2, Ser48) are unique serines in CML14 that represent phospho-sites found via global phosphoproteomic analyses (Fig. 1A) (Willems *et al*., 2019). In addition, Ser2 in CML14 has been identified as an acetylation site (Willems *et al*., 2019). Although speculative, it is possible that differences in post-translational modification could contribute to the lack of redundancy between these CMLs, perhaps by altering target or IQ-domain binding specificity or affinity *in vivo*.

Although CML13 and CML14 are almost identical at the primary sequence level, differing by only eight residues, our data indicate that they are not fully redundant *in vivo*, as reduction of either transcript resulted in strong phenotypic consequences. There are several possible explanations that may account for this. If both CML13 and CML14 are required to interact with one or more identical targets in unison for proper function, the inhibition of either CML could lead to the same deleterious phenotype. For example, CML13 and CML14 may have different, specific targets among the various IQ-domain proteins that are involved in the same pathways, and/or they may bind to distinct IQ domains on some of their common targets and thus both CMLs may be required for full function of certain targets. It should be noted that CAMTAs, IQDs, and myosins, all possess between two and six IQ domains, and considerable sequence variation is found among the IQ domains of any given family member. An example of this can be seen for some myosins in other organisms, where CaM serves as the essential light chain on a specific IQ domain but other, CaM-related proteins function as a regulatory light chain on an adjacent IQ domain (Heissler and Sellers, 2014). For example, if a given myosin, with multiple IQ domains, simultaneously requires CaM, CML13, and CML14, as light chains for motility, loss on any of these interactors would be expected to yield a similar phenotype. This is consistent with the fact that in our study, comparable phenotypes were observed for either *CML13* or *CML14* hpRNAi lines. The simplest hypothesis is that CML13 and CML14 interact with the same (or overlapping) IQ-domain targets as part of a larger complex. This is supported by binding assays showing that CML13 and CML14 have broadly overlapping targets among CAMTAs, IQDs, and myosins (Teresinski *et al*., 2023). Given that CaM also interacts with these same IQ-domain targets, an important area for future investigation will be to distinguish between the roles of CaM, CML13, and CML14 in the regulation of target function and, ultimately, how this activity contributes to plant development.

## Materials and Methods

### Plasmid Construction

The dexamethasone (Dex)-inducible RNA interference (RNAi) plasmids were generated using a golden gate assembly system (Binder *et al*., 2014; Chiasson *et al*., 2019). Level-III plant binary plasmids were built using pre-assembled LII transcription units. LII units consisted of a kanamycin selection cassette, an inducible RNAi hairpin (RNAi-hp) expression unit, and the artificial GR-LhG4 transcriptional activator. RNAi fragments from the 3’UTRs of *CML13* (At1g12310; 249 bp) and *CML14* (At1g62820; 236 bp) were amplified by PCR to generate LI C-D entry clones (Supplementary Table S1). Each fragment was then combined with a LI *AtWRKY33* intron part (Maekawa *et al*., 2008) in a custom LII backbone containing the pOp6 35Smin promoter (Craft *et al*., 2005) and the 35S terminator. GR-LhG4 consists of the lac repressor DNA binding domain, a GAL4 activation domain, and the ligand-binding domain of the rat glucocorticoid receptor (Craft *et al*., 2005). Constitutive *GR-LhG4* expression was driven by the At*UBQ10* promoter. The pOp6 35Smin promoter and the GR-LhG4_BD transcriptional activator were amplified from pSW180a-pOp6 and pSW610-GR-LhG4_BD respectively (Schürholz *et al*., 2018).

### Protein Expression and Purification

Recombinant CML13, CML14, and CaM were purified as outlined previously (Teresinski *et al*., 2023). Briefly, the open-reading frames of *CML13*, *CML14*, or *CaM7* (At3g43810, a conserved isoform of CaM) were subcloned into pET21a expression vectors (Novagen) as His-tagged fusion proteins and were expressed in BL21 CPRL *E. coli* cells (Novagen) as described (Dobney et al., 2009). Recombinant CML13 and CML14 were purified via Ni-NTA His60 (Takara Biosciences.) and CaM7 using phenyl-sepharose (Sigma Aldrich) as described (Ogunrinde *et al*., 2017). Protein purity was assessed by SDS-PAGE and concentrations were determined by the Bradford Assay and absorbance at 280 nm.

### Plant Material and Growth Conditions

*Arabidopsis thaliana* (Arabidopsis) accession Columbia-0 (WT) seeds were sown into Sunshine mix #1 or 0.5X MS (Murashige and Skoog, 1962) basal salt media agar plates and stratified at 4°C for 3-5 days in the dark before transfer to a growth chamber under long-day conditions unless otherwise stated (Sun Gro Horticulture Canada Ltd., and Conviron MTR30; 18h photoperiod, 22°C, ∼150 μmol m^−2^ s^−1^). Plants were watered and fertilized as needed. Transgenic Arabidopsis were generated by the floral dip method (Zhang *et al*., 2006; Mireault *et al*., 2014) and transformants were selected using kanamycin (50 µg/mL) and 0.5X MS agar plates. Segregation analysis was conducted to select transgenic lines carrying a single-transgene insertion and these lines were progressed to homozygosity for RT-qPCR and phenotypic analyses. Seeds for WT plants were always sterilized, sown, and grown alongside transgenic lines and all genotypes were cultivated and maintained under identical conditions.

### Plant Phenotyping

For plate assays, Arabidopsis seeds were sown or transplanted onto 0.5X MS agar medium with or without 5 µM dexamethasone as noted in the figure legends. Cotyledon greening was tabulated as a percentage of green:yellow cotyledons on a given plate (Liu *et al*., 2019) and root lengths were measured as the growth of the primary root per day after transplanting. Root lengths were marked on the petri dishes for four days before the plates were imaged and analyzed using ImageJ software (Schneider *et al*., 2012) for root length per day measured in cm. For survival assays, Arabidopsis Col-0 and mutant seeds were sown onto petri plates containing 25 mL of ½ MS + 0.8% Agar and allowed to grow for seven days after stratification. Plates were then supplemented with 5 mL of milliQ water or a 40 µM Dex solution for control or treatment groups, respectively and allowed to grow for another seven days. Percent survival was scored by the ratio of seedlings with:without yellowing or white tissue.

Plants grown in soil were sprayed with a solution of 0.05% Silwett L-77 (Lehle Seeds Inc) with or without 20 µM dexamethasone every 72 hours for the extent of the treatment course as indicated in the figure legends. All phenotyping was performed according to the standard protocols of Boyes *et al*. (2001) unless otherwise stated.

### Chlorophyll Content

Seedlings used in the percent greening assay were removed from the agar medium and weighed on an analytical balance to record their fresh weights before submerging them in 1 mL of 80% acetone overnight in the dark. Tubes were then centrifuged, and the supernatant was decanted into a cuvette. Chlorophyll content was calculated as described (Remy and Duque, 2016) using the following equation: (18.71*Abs_647nm_+ 7.15*Abs_660nm_) / fresh weight.

### RT-qPCR

Plants were sown onto 0.5X MS media as described above and grown for 7 days before harvesting tissue and flash freezing in liquid nitrogen. Frozen plant tissue was ground to a powder and RNA was extracted using Monarch ® Total RNA Miniprep kits (NEB), RNA integrity and concentration were assessed by agarose gels and a Nanodrop spectrophotometer (Thermo Fisher Scientific). RT-qPCR was performed using a Luna ® Universal One-Step RT-qPCR kit (NEB) in a CFX96 qPCR thermocycler (BioRad). All data were normalized to the reference gene Actin2 as previously described (Graeber *et al*., 2011; Zhuang *et al*., 2013), Cq values (the cycle number at which the threshold fluorescence level was reached in a qPCR reaction) were extracted, and relative expression was calculated in Excel (Microsoft) before exporting to GraphPad Prism 9 for statistical analysis.

### Differential Scanning Fluorimetry

Samples of pure, recombinant CML13 or CML14 at concentrations of 15, 20, and 25 µM were mixed with SYPRO Orange (Thermo Fisher Scientific) at 5X concentration and adjusted to 1 mM CaCl_2_ or EGTA as noted in the figure legends. The temperature ramp rate was 1°C/20 s and protein unfolding was monitored by the increase in RFUs produced every 10 s in a CFX Opus 96 System (BioRad). The average trace of 6 replications (2 at each concentration) was plotted for each CML and condition. The T_m_ was determined by identifying the global minima of dRFU/dT of each rep. Data are presented as mean +/− SEM.

### Steady State ANS Fluorimetry

The hydrophobic properties of recombinant CaM7, CML13, and CML14 were monitored using ANS (8-anilinonapthalene-1-sulfonic acid) fluorescence emission spectroscopy using an excitation wavelength of 380 nm, and scanning emission spectra 430–600 nm, as described (Ogunrinde *et al*., 2017). Purified, recombinant proteins were dialyzed overnight against one L of ANS buffer (10 mM Tris-Cl, pH 7.5, containing 1 mM DTT, and 100 mM KCl). The fluorescence emission of 20 μM CML13 and 20 μM CML14 was recorded using 250 μM ANS in ANS buffer in the presence or absence of calcium as noted in figure legends. Background ANS fluorescence in each buffer was subtracted and the average trace of three replicates was plotted.

### Data Analysis

Data graphs and statistical analysis were performed using GraphPad Prism 9. Unless otherwise stated a 2-way ANOVA with Sidak’s test for multiple comparisons was performed and, ns is not significantly different, p-value * < .1, ** < .01, *** < .001, **** < .0001.

## Supplementary Data Information

Supplementary Table S1. List of oligonucleotide primers used in PCR.

**Supplementary Figure 1.**
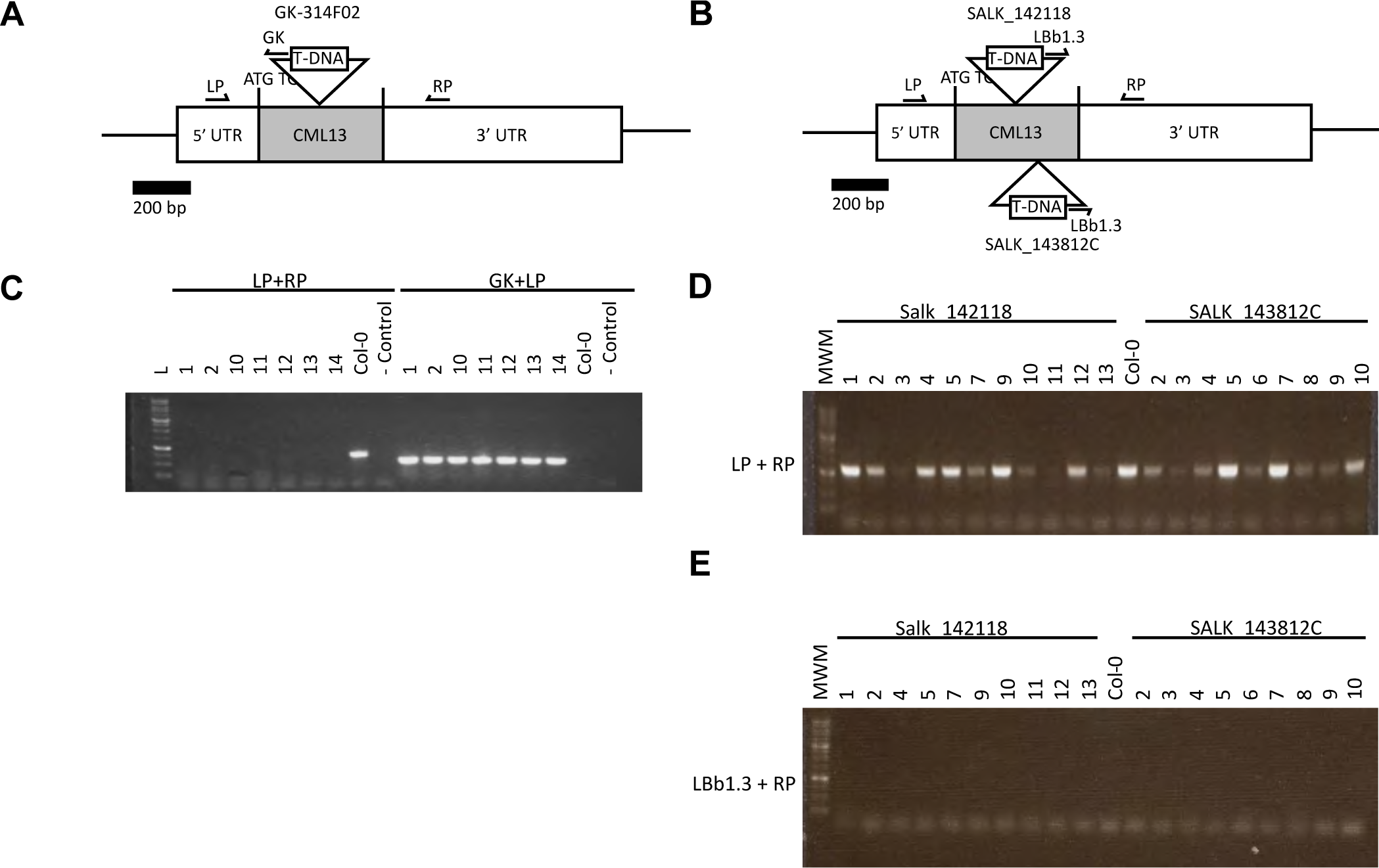
Schematic diagram showing the gene structure and relative, reported T-DNA positions in the exon of (A) CML13 (At1g12310) in GABI-Kat line GK-314F02 and (B) CML14 (At1g62820) in lines SALK_142118 and SALK_143812C. Both At1g12310 and At1g62820 are intron-less genes, and the single exons are shaded in grey. PCR primers used to screen genomic DNA from Col-0 and mutant lines are indicated by black arrows LP, GK, LBb1.3, and RP and are described in Supplementary Table 1. (C) PCR amplification of the wild type (Col-0) or GK-314F02 CML13 allele using combinations of forward (LP), reverse (RP), and GK primers on genomic DNA from seven sibling seedlings (numbered above panel) homozygous for the T-DNA insertion. Control lanes lacked genomic DNA template. PCR analysis of wild type (Col-0) or two putative T-DNA insertion lines for CML14 allele using combinations of (D) forward (LP) and reverse (RP) primers or (E) LBb1.3 and RP primers on genomic DNA from random sibling seedlings.

**Supplementary Figure 2.**
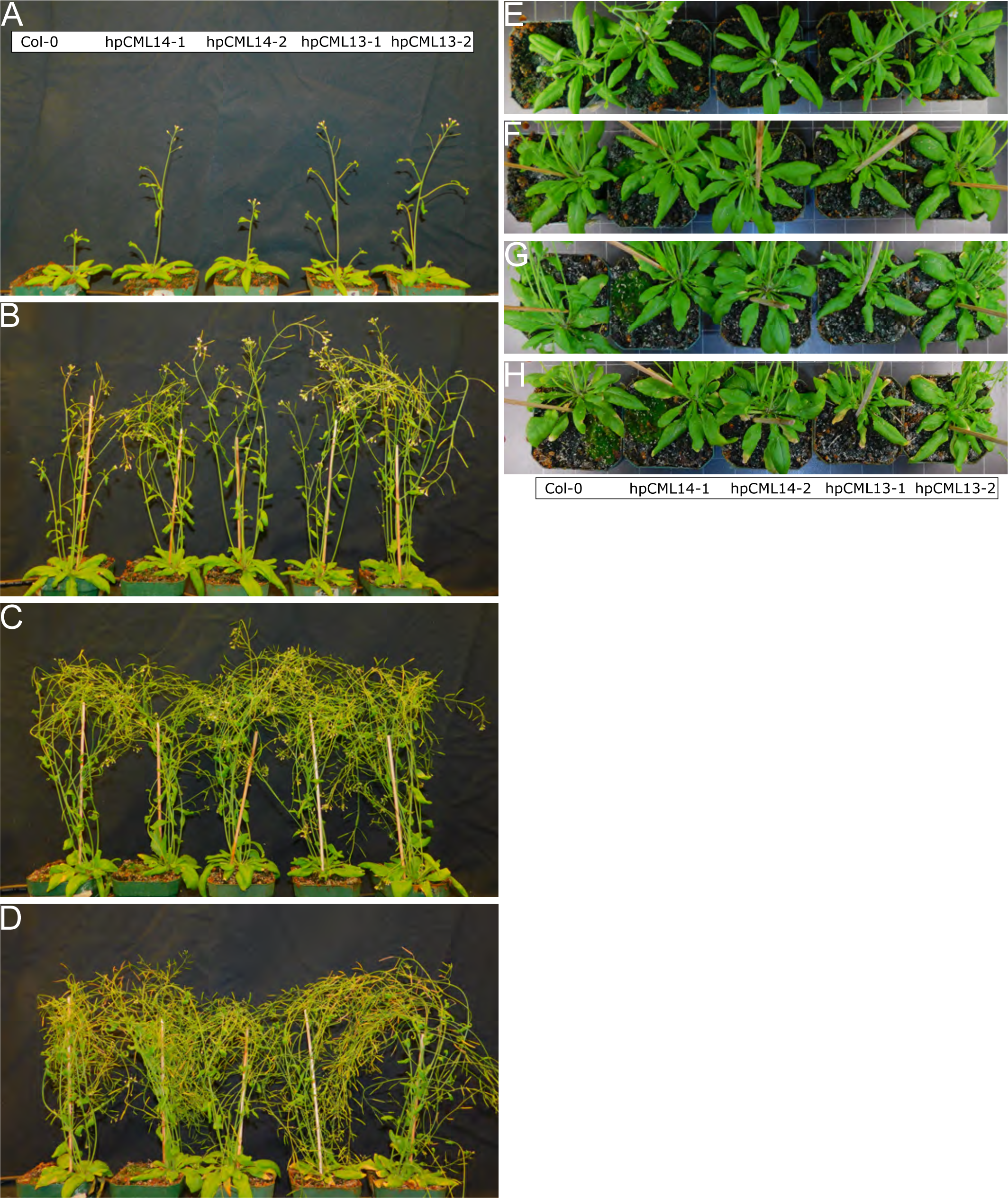
Developmental time-course showing representative side-view (A-D) and overhead (E-H) images of plant growth for WT and hpCML RNAi untreated plants at (A, E) 4 weeks, (B, F) 5 weeks, (C, G) 6 weeks, and (D, H) 7 weeks of growth. Genotypes are indicated on or below the panels. Each panel shows the identical group of plants over the 7-week period. Experiments were done in triplicate with a minimum of three technical replicates.

**Supplementary Figure 3.**
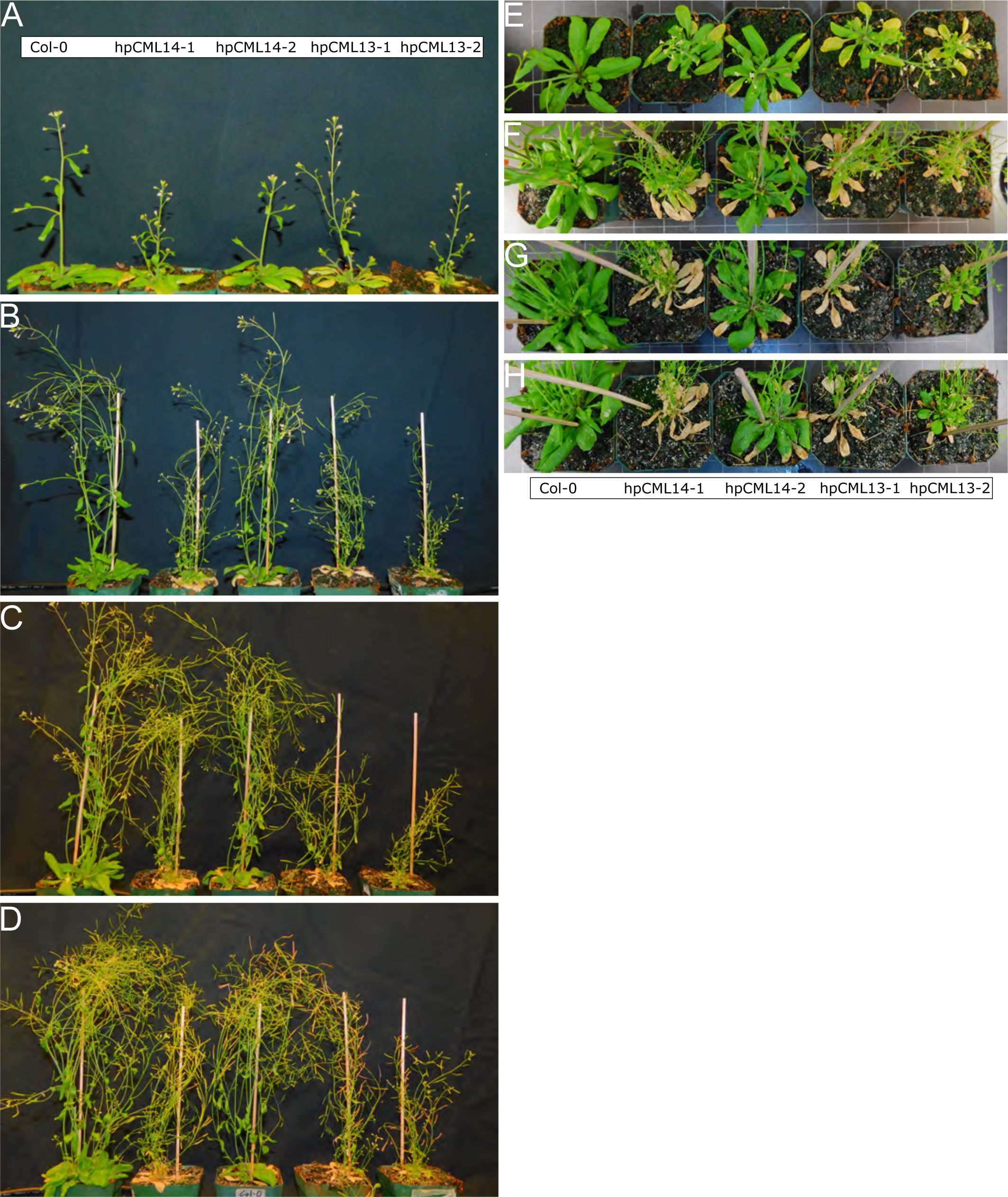
Developmental time-course showing representative side-view (A-D) and overhead (E-H) images of plant development of WT and hpCML RNAi plants sprayed with a solution of 20 uM Dex every 72h starting at 3 weeks of growth and imaged subsequently at (A, E) 4 weeks, (B, F) 5 weeks, (C, G) 6 weeks, or (D, H) 7 weeks of growth. Genotypes are indicated on or below the panels. Each panel shows the identical group of plants over the 7-week period. Experiments were done in triplicate with a minimum of three technical replicates.

**Supplementary Figure 4.**
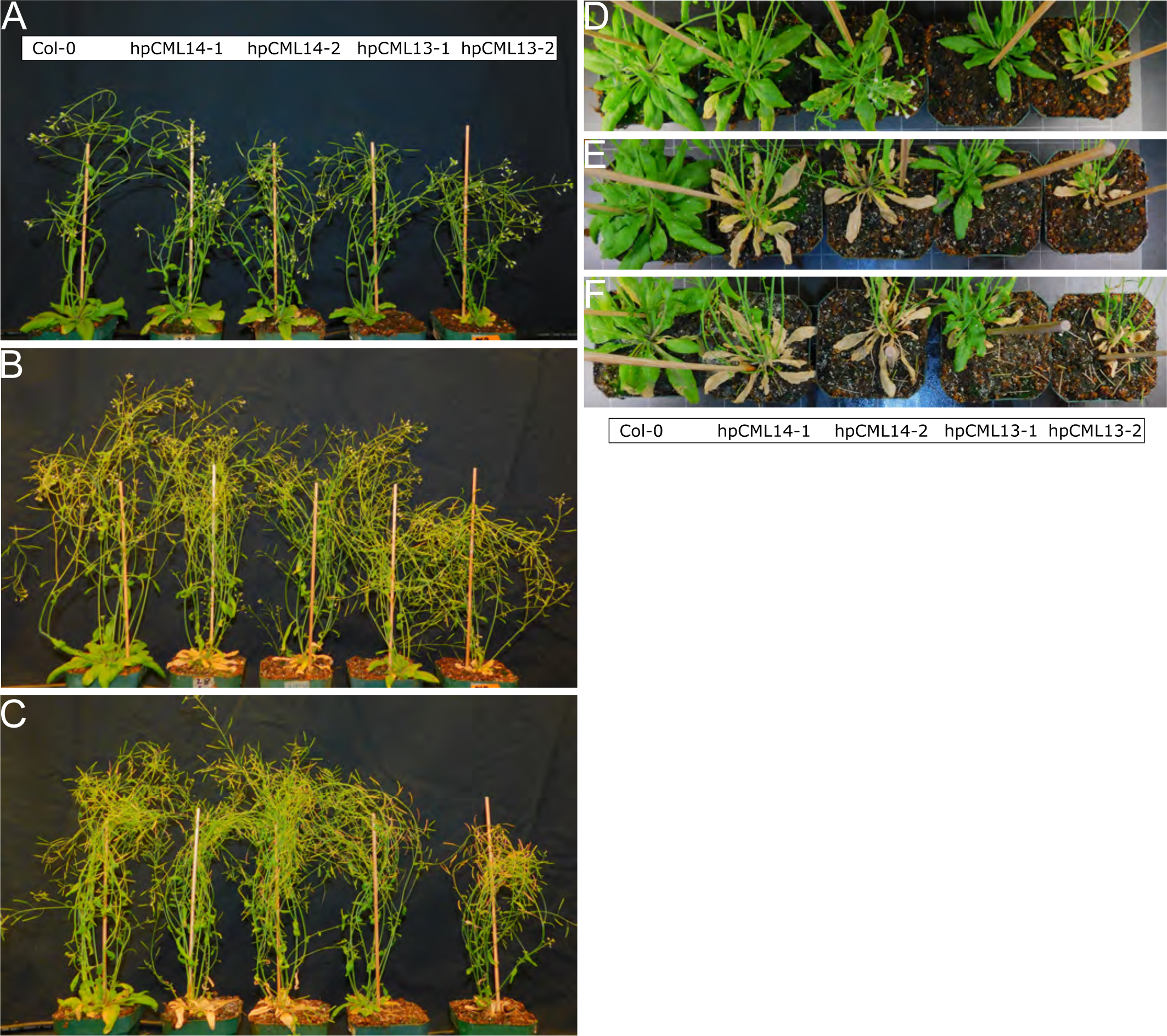
Developmental time-course showing representative side-view (A-D) and overhead (E-H) images of plant development of WT and hpCML RNAi plants sprayed with a solution of 20 uM Dex every 72h starting at 4 weeks of growth and imaged subsequently at (A, D) 5 weeks, (B, E) 6 weeks, (C, F) 7 weeks of growth. Genotypes are indicated on or below the panels. Each panel shows the identical group of plants over the 7-week period. Experiments were done in triplicate with a minimum of three technical replicates.

**Supplementary Figure 5.**
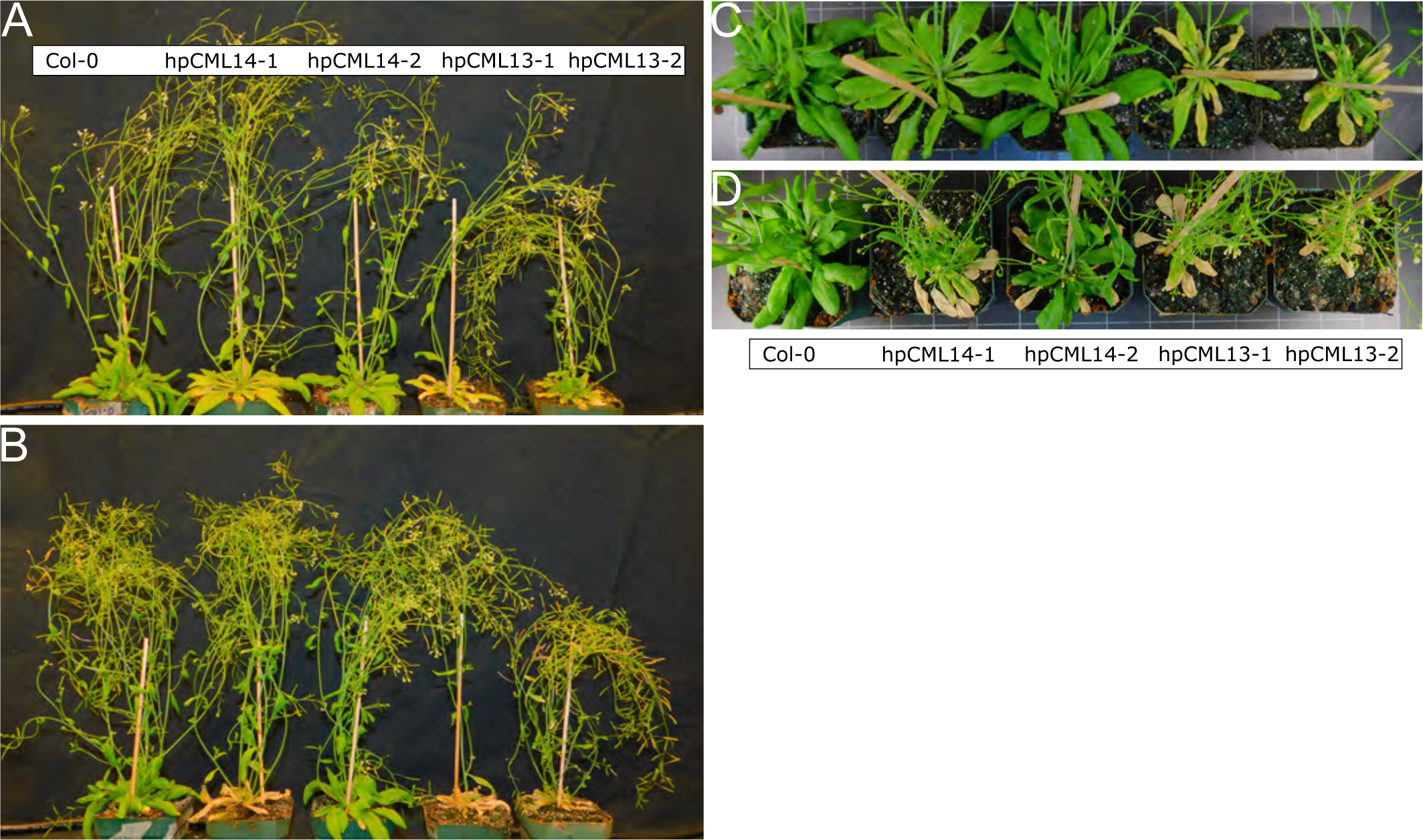
Developmental time-course showing representative side-view (A-D) and overhead (E-H) images of plant development of WT and hpCML RNAi plants sprayed with a solution of 20 uM Dex every 72h starting at 5 weeks of growth and imaged subsequently at (A, C) 6 weeks, (B, D) 7 weeks of growth. Genotypes are indicated on or below the panels. Each panel shows the identical group of plants over the 7-week period. Experiments were done in triplicate with a minimum of three technical replicates.

**Supplementary Figure 6.**
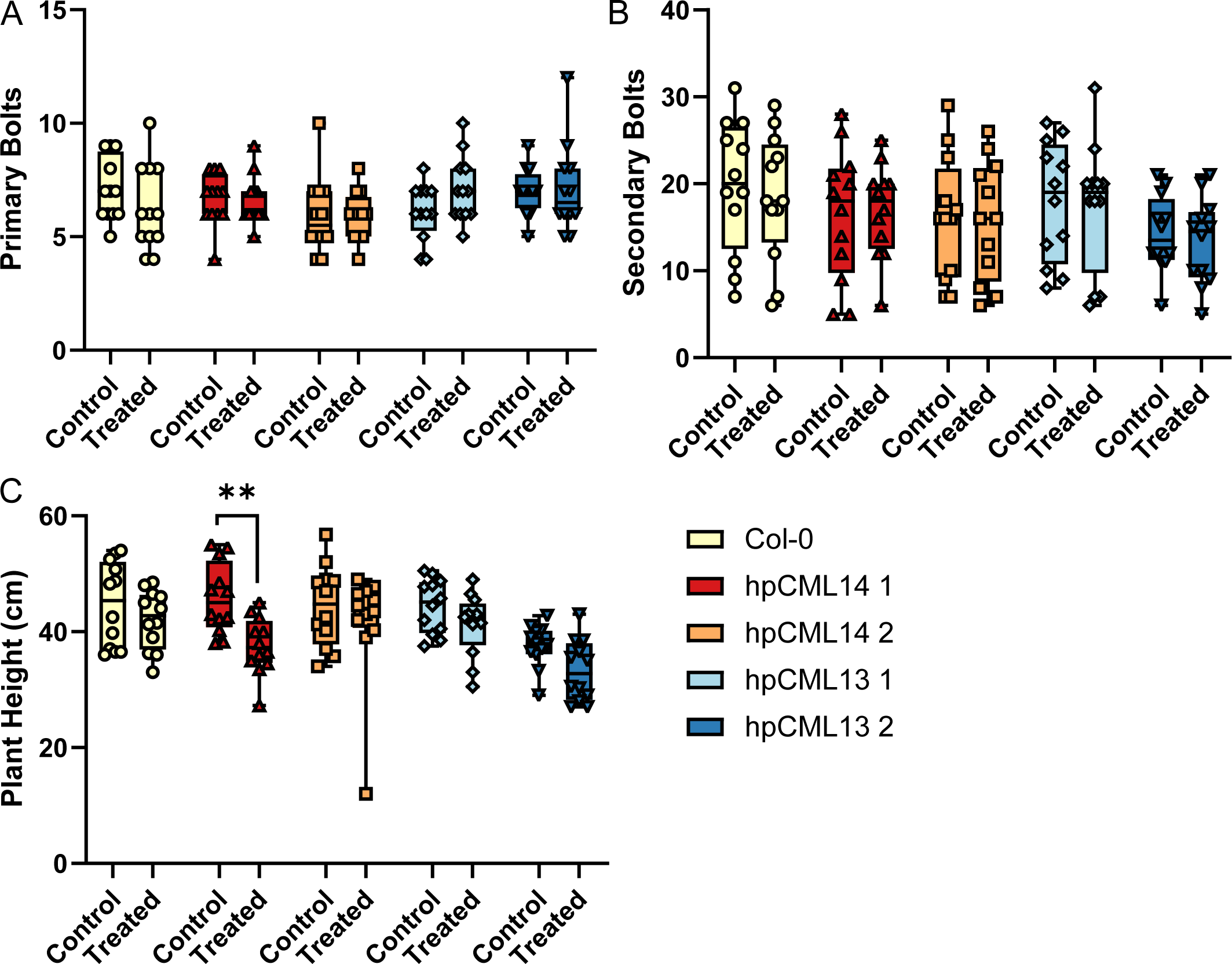
Dex treatment at later stages of development does not have a strong impact on hpCML RNAi growth or morphology. Quantitative analysis of (A) number of primary bolts, (B) secondary bolts, and (C) plant height for WT (Col-0) and hpCML RNAi plants grown without Dex (control) or sprayed with 20 uM Dex every 72h starting at 4 weeks of growth (same plants as were used for the silique measurements). Box plot of silique length for control and Dex-treated plants. Points are the lengths of 30 siliques in each control or 20 µM Dex-treated groups. (2-way ANOVA with Sidak’s test for multiple comparisons, ns is not significantly different, p-value * < .1, ** < .01, *** < .001, **** < .0001).

## Data Availability

All data supporting the findings of this study are available in this paper and in its supplementary data published online.

## Author Contributions

WS, KS, HT, BH, DC, designed and planned the study; KS, HT, BH, DC, performed the experiments and analyses; all authors contributed to the writing, editing and approval of the manuscript.

## References

Astegno, A., Bonza, M.C., Vallone, R., and La Verde, V. (2017) Arabidopsis calmodulin-like protein CML36 is a calcium (Ca^2+^) sensor that interacts with the plasma membrane Ca^2+^-ATPase isoform ACA8 and stimulates its activity. Journal of Biological Chemistry. 292: 15049–15061.

Bac-Molenaar, J.A., Fradin, E.F., Becker, F.F.M., Rienstra, J.A., van der Schoot, J., Vreugdenhil, D., and Keurentjes, J.J.B. (2015) Genome-Wide Association Mapping of Fertility Reduction upon Heat Stress Reveals Developmental Stage-Specific QTLs in *Arabidopsis thaliana*. The Plant Cell. 27: 1857–1874.

Bender, K.W., and Snedden, W.A. (2013) Calmodulin-related proteins step out from the shadow of their namesake. Plant Physiology. 163: 486–495.

Bender, K.W., Rosenbaum, D.M., Vanderbeld, B., Ubaid, M., and Snedden, W.A. (2013) The Arabidopsis calmodulin-like protein, CML39, functions during early seedling establishment. The Plant Journal. 76: 634–647.

Berridge, M.J., Lipp, P., and Bootman, M.D. (2000) The versatility and universality of calcium signalling. Nature Reviews Molecular Cell Biology. 1: 11–21.

Binder, A., Lambert, J., Morbitzer, R., Popp, C., Ott, T., Lahaye, T., and Parniske, M. (2014) A modular plasmid assembly kit for multigene expression, gene silencing and silencing rescue in plants. PloS One. 9: e88218.

Boyes, D.C., Zayed, A.M., Ascenzi, R., McCaskill, A.J., Hoffman, N.E., Davis, K.R., and Görlach, J. (2001) Growth Stage–Based Phenotypic Analysis of Arabidopsis: A Model for High Throughput Functional Genomics in Plants. The Plant Cell. 13: 1499–1510.

Brueggeman, J.M., Windham, I.A., and Nebenführ, A. (2022) Nuclear movement in growing Arabidopsis root hairs involves both actin filaments and microtubules. Journal of Experimental Botany. 73: 5388–5399.

Buschmann, H. (2018) Myosin XI motors: back on the scene at the division machine. Journal of Experimental Botany. 69: 2863–2866.

Chiasson, D., Giménez-Oya, V., Bircheneder, M., Bachmaier, S., Studtrucker, T., Ryan, J., Sollweck, K., Leonhardt, H., Boshart, M., Dietrich, P., and Parniske, M. (2019) A unified multi-kingdom Golden Gate cloning platform. Scientific Reports. 9: 10131.

Clapham, D.E. (2007) Calcium signaling. Cell. 131: 1047–1058.

Craft, J., Samalova, M., Baroux, C., Townley, H., Martinez, A., Jepson, I., Tsiantis, M., and Moore, I. (2005) New pOp/LhG4 vectors for stringent glucocorticoid-dependent transgene expression in Arabidopsis. The Plant Journal. 41: 899–918.

DeFalco, T.A., Bender, K.W., and Snedden, W.A. (2009) Breaking the code: Ca2+ sensors in plant signaling. Biochemical Journal. 425: 27–40.

Dobney, S., Chiasson, D., Lam, P., Smith, S.P., and Snedden, W.A. (2009) The calmodulin-related calcium sensor CML42 plays a role in trichome branching. Journal of Biological Chemistry. 284: 31647–31657.

Edel, K.H., Marchadier, E., Brownlee, C., Kudla, J., and Hetherington, A.M. (2017) The Evolution of Calcium-Based Signalling in Plants. Current Biology. 27: R667–R679.

Finkler, A., Ashery-Padan, R., and Fromm, H. (2007) CAMTAs: calmodulin-binding transcription activators from plants to human. FEBS Letters. 581: 3893–3898.

Fischer, C., DeFalco, T.A., Karia, P., Snedden, W.A., Moeder, W., Yoshioka, K., and Dietrich, P. (2017) Calmodulin as a Ca^2+^-Sensing Subunit of Arabidopsis Cyclic Nucleotide-Gated Channel Complexes. Plant and Cell Physiology. 58: 1208–1221.

Golomb, L., Abu-Abied, M., Belausov, E., and Sadot, E. (2008) Different subcellular localizations and functions of Arabidopsis myosin VIII. BMC Plant Biology, 8(3) doi: 10.1186/1471-2229-8-3.

Graeber, K., Linkies, A., Wood, A.T.A., and Leubner-Metzger, G. (2011) A guideline to family-wide comparative state-of-the-art quantitative RT-PCR analysis exemplified with a Brassicaceae cross-species seed germination case study. The Plant Cell. 23: 2045–2063.

Haraguchi, T., Duan, Z., Tamanaha, M., Ito, K., and Tominaga, M. (2019) Diversity of Plant Actin–Myosin Systems. In: Sahi, V.P., and Baluška, F. (Eds.), The Cytoskeleton: Diverse Roles in a Plant’s Life (pp. 49–61) Cham: Springer International Publishing.

Heissler, S.M., and Sellers, J.R. (2014) Myosin light chains: Teaching old dogs new tricks. Bioarchitecture. 4: 169–188.

Iqbal, Z., Shariq Iqbal, M., Singh, S.P., and Buaboocha, T. (2020) Ca2+/Calmodulin Complex Triggers CAMTA Transcriptional Machinery Under Stress in Plants: Signaling Cascade and Molecular Regulation. Frontiers in Plant Science. 11: 598327.

Jeon, Y., Park, Y.-J., Cho, H.K., Jung, H.J., Ahn, T.-K., Kang, H., and Pai, H.-S. (2015) The nucleolar GTPase nucleostemin-like 1 plays a role in plant growth and senescence by modulating ribosome biogenesis. Journal of Experimental Botany. 66: 6297–6310.

Khan, B.R., and Zolman, B.K. (2010) pex5 Mutants that differentially disrupt PTS1 and PTS2 peroxisomal matrix protein import in Arabidopsis. Plant Physiology. 154: 1602–1615.

Kim, M.C., Chung, W.S., Yun, D.-J., and Cho, M.J. (2009) Calcium and calmodulin-mediated regulation of gene expression in plants. Molecular Plant. 2: 13–21.

Kumari, P., Dahiya, P., Livanos. P., Zergiebel. L., Kölling. M., Poeschl, Y., Stamm, G., Hermann, A., Abel, S., Müller, S., and Bürstenbinder, K. (2021) IQ67 DOMAIN proteins facilitate preprophase band formation and division-plane orientation. Nature Plants. 7: 739– 747.

Langelaan, D.N., Liburd, J., Yang, Y., Miller, E., Chitayat, S., Crawley, S.W., Côté, G..P, and Smith, SP. (2016) Structure of the single-lobe myosin light chain c in complex with the light chain-binding domains of myosin-1c provides insights into divergent IQ motif recognition. The Journal of Biological Chemistry. 291: 19607–19617.

La Verde, V., Dominici, P., and Astegno, A. (2018a) Towards Understanding Plant Calcium Signaling through Calmodulin-Like Proteins: A Biochemical and Structural Perspective. International Journal of Molecular Sciences. 19: 1331.

La Verde, V., Trande, M., D’Onofrio, M., Dominici, P., and Astegno, A. (2018b) Binding of calcium and target peptide to calmodulin-like protein CML19, the centrin 2 of *Arabidopsis thaliana*. International Journal of Biological Macromolecules. 108: 1289–1299.

Li, L., Nelson C.J., Trösch, J., Castleden, I., Huang, S., and Millar, A.H. (2017) Protein Degradation Rate in *Arabidopsis thaliana* Leaf Growth and Development. The Plant Cell. 29: 207–228.

Liu, H., Guo, S., Lu, M., Zhang, Y., Li, J., Wang, W., et al. (2019) Biosynthesis of DHGA12 and its roles in Arabidopsis seedling establishment. Nature Communications. 10: 1768.

Liu, S., and Yoder, J.I. (2016) Chemical induction of hairpin RNAi molecules to silence vital genes in plant roots. Scientific Reports. 6: 37711.

Lokdarshi, A., Conner, W.C., McClintock, C., Li, T., and Roberts, D.M. (2016) Arabidopsis CML38, a Calcium Sensor That Localizes to Ribonucleoprotein Complexes under Hypoxia Stress. Plant Physiology. 170: 1046–1059.

Luan, S., and Wang, C. (2021) Calcium Signaling Mechanisms Across Kingdoms. Annual Review of Cell and Developmental Biology. 37: 311–340.

Madison, S.L., Buchanan, M.L., Glass, J.D., McClain, T.F., Park, E., and Nebenführ, A. (2015) Class XI Myosins Move Specific Organelles in Pollen Tubes and Are Required for Normal Fertility and Pollen Tube Growth in Arabidopsis. Plant Physiology. 169: 1946–1960.

Maekawa, T., Kusakabe, M., Shimoda, Y., Sato, S., Tabata, S., Murooka, Y., and Hayashi, M. (2008) Polyubiquitin promoter-based binary vectors for overexpression and gene silencing in Lotus japonicus. Molecular Plant-Microbe Interactions. 21: 375–382.

McCormack, E., and Braam, J. (2003) Calmodulins and related potential calcium sensors of Arabidopsis. The New Phytologist. 159: 585–598.

Midhat, U., Ting, M.K.Y., Teresinski, H.J., and Snedden, W.A. (2018) The calmodulin-like protein, CML39, is involved in regulating seed development, germination, and fruit development in Arabidopsis. Plant Molecular Biology. 96: 375–392.

Mireault, C., Paris, L-E., and Germain, H. (2014) Enhancement of the Arabidopsis floral dip method with XIAMETER OFX-0309 as an alternative to Silwet L-77 surfactant. Botany. 92: 523–525.

Mitra, D., Klemm, S., Kumari, P., Quegwer, J., Möller, B., Poeschl, Y., et al. (2019) Microtubule-associated protein IQ67 DOMAIN5 regulates the morphogenesis of leaf pavement cells in *Arabidopsis thaliana*. Journal of Experimental Botany. 70: 529–543.

Murashige, T., and Skoog, F. (1962) A Revised Medium for Rapid Growth and Bio Assays with Tobacco Tissue Cultures. Physiologia Plantarum. 15: 473–497.

Nie, H., Zhao, C., Wu, G., Wu, Y., Chen, Y., and Tang, D. (2012) SR1, a calmodulin-binding transcription factor, modulates plant defense and ethylene-induced senescence by directly regulating NDR1 and EIN3. Plant Physiology. 158: 1847–1859.

Ogunrinde, A., Munro, K., Davidson, A., Ubaid, M., and Snedden, W.A. (2017) Calmodulin-Like Proteins, CML15 and CML16 Possess Biochemical Properties Distinct from Calmodulin and Show Non-overlapping Tissue Expression Patterns. Frontiers in Plant Science. 8: 2175.

Peremyslov, V.V., Prokhnevsky, A.I., and Dolja, V.V. (2010) Class XI myosins are required for development, cell expansion, and F-Actin organization in Arabidopsis. The Plant Cell. 22: 1883–1897.

Remy, E., and Duque, P. (2016) Assessing Tolerance to Heavy-Metal Stress in Arabidopsis thaliana Seedlings. Methods Mol Biol.1398: 197–208.

Schneider, C.A., Rasband, W.S., and Eliceiri KW. (2012) NIH Image to ImageJ: 25 years of image analysis. Nature Methods. 9: 671–675.

Scholz, S.S., Vadassery, J., Heyer, M., Reichelt, M., Bender, K.W., Snedden, W.A. et al. (2014) Mutation of the Arabidopsis calmodulin-like protein CML37 deregulates the jasmonate pathway and enhances susceptibility to herbivory. Molecular Plant. 7: 1712–1726.

Schubert, J., Li, Y., Mendes, M.A., Fei, D., Dickinson, H., Moore, I., and Baroux, C. (2022) A procedure for Dex-induced gene transactivation in Arabidopsis ovules. Plant Methods. 18: 41.

Schürholz, A.K., López-Salmerón, V., Li, Z., Forner, J., Wenzl, C., Gaillochet, C., et al. (2018) A Comprehensive Toolkit for Inducible, Cell Type-Specific Gene Expression in Arabidopsis. Plant Physiology. 178: 40–53.

Shkolnik, D., Finkler, A., Pasmanik-Chor, M., and Fromm, H. (2019) CALMODULIN-BINDING TRANSCRIPTION ACTIVATOR 6: A Key Regulator of Na+ Homeostasis during Germination. Plant Physiology. 180: 1101–1118.

Symonds, K., Teresinski, H.J., Hau, B., Sadot, E., Dwivedi, V., Belausov, E., Bar-Sinai S, Tominaga, M., Haraguchi, T., Ito, K., and Snedden, W.A. (2023) Functional Characterization of Calmodulin-like Proteins, CML13 and CML14, as Novel Light Chains of Arabidopsis Class VIII Myosins, Biorxiv preprint, doi: 10.1101/2023.05.12.540561

Tang. R.J., Wang, C., Li, K., and Luan, S. (2020) The CBL-CIPK Calcium Signaling Network: Unified Paradigm from 20 Years of Discoveries. Trends Plant Science. 25: 604–617.

Teresinski, H.J., Hau, B., Symonds, K., Kilburn, R., Munro, K.A., Doner, N.M., Mullen, R., Li, V.H., and Snedden, W.A. (2023) Arabidopsis calmodulin-like proteins CML13 and CML14 interact with proteins that have IQ domains. Plant Cell Environ. doi: 10.1111/pce.146 Epub ahead of print.

Tian, X., Wang, X., and Li Y. (2021) Myosin XI-B is involved in the transport of vesicles and organelles in pollen tubes of *Arabidopsis thaliana*. The Plant Journal. 108: 1145–1161.

Tsai, Y-C., Koo, Y., Delk, N.A., Gehl, B., and Braam, J. (2013) Calmodulin-related CML24 interacts with ATG4b and affects autophagy progression in Arabidopsis. The Plant Journal. 73: 325–335.

Vadassery, J., Reichelt, M., Hause, B., Gershenzon, J., Boland, W., and Mithöfer, A. (2012) CML42-mediated calcium signaling coordinates responses to Spodoptera herbivory and abiotic stresses in Arabidopsis. Plant Physiology. 159: 1159–1175.

Vallone, R., La Verde, V., D’Onofrio, M., Giorgetti, A., Dominici, P., and Astegno, A. (2016) Metal binding affinity and structural properties of calmodulin-like protein 14 from *Arabidopsis thaliana*. Protein Science. 25: 1461–1471.

Vanderbeld, B., and Snedden WA. (2007) Developmental and stimulus-induced expression patterns of Arabidopsis calmodulin-like genes CML37, CML38 and CML Plant Molecular Biology. 64: 683–697.

Van Leene, J., Eeckhout, D., Gadeyne, A., Matthijs, C., Han, C., De Winne, N., et al. (2022) Mapping of the plant SnRK1 kinase signalling network reveals a key regulatory role for the class II T6P synthase-like proteins. Nature Plants. 8: 1245–1261.

Wang, M., Herrmann, C.J., Simonovic, M., Szklarczyk, D., and Mering, C. (2015) Version 4.0 of PaxDb: Protein abundance data, integrated across model organisms, tissues, and cell-lines. Proteomics. 15: 3163–3168.

Willems, P., Horne, A., Van Parys, T., Goormachtig, S., De Smet, I., Botzki, A., Van Breusegem, F., and Gevaert, K. (2019) The Plant PTM Viewer, a central resource for exploring plant protein modifications. The Plant Journal. 99.: 752–762.

Yip Delormel, T., and Boudsocq, M. (2019) Properties and functions of calcium-dependent protein kinases and their relatives in Arabidopsis thaliana. New Phytologist. 224: 585–604.

Zhang, X., Henriques, R., Lin, S-S., Niu, Q-W., and Chua, N-H. (2006) Agrobacterium-mediated transformation of *Arabidopsis thaliana* using the floral dip method. Nature Protocols. 1: 641–646.

Zhuang, X., Wang, H., Lam, S.K., Gao, C., Wang, X., Cai, Y., and Jiang, L. (2013) A BAR-domain protein SH3P2, which binds to phosphatidylinositol 3-phosphate and ATG8, regulates autophagosome formation in Arabidopsis. The Plant Cell. 25: 4596–4615.

Zhu, X., Dunand, C., Snedden, W., and Galaud, J-P. (2015) CaM and CML emergence in the green lineage. Trends in Plant Science. 20: 483–489.

Zhu, X., Perez, M., Aldon, D., and Galaud, J-P. (2017) Respective contribution of CML8 and CML9, two Arabidopsis calmodulin-like proteins, to plant stress responses. Plant Signaling and Behavior. 12: e1322246.

